# The DNA damage response in myogenic C2C7 cells depends on the characteristics of ionizing particles

**DOI:** 10.1101/2024.06.24.600399

**Authors:** Haser H. Sutcu, Arthur Thomas--Joyeux, Mikaël Cardot-Martin, Delphine Dugué, François Vianna, Yann Perrot, Mohamed A. Benadjaoud, Marc Benderitter, Céline Baldeyron

**Author notes:** Université Paris Cité, Inserm, CEA, Stabilité Génétique Cellules Souches et Radiations/iRCM, F-92265, Fontenay-aux-Roses, France; Université Paris-Saclay, Inserm, CEA, Stabilité Génétique Cellules Souches et Radiations/iRCM, F-92265, Fontenay-aux-Roses, France. **Co-corresponding authors:** Céline Baldeyron,; Haser Hasan Sutcu.

## Abstract

DNA integrity and stability are vital for proper cellular activity. Nevertheless, to treat cancer patients, DNA is the main target for inducing tumoral cell death. Nowadays, cancer treatment is improving by the development of new technologies, protocols and strategies. Amongst them, the charged particle radiotherapies are becoming prevalent. However, tumor-neighboring healthy tissues are still exposed to ionizing radiations (IR) and subject to late side effects. Skeletal muscle is one of those tissues most likely to be affected. To decipher the DNA damage response (DDR) of skeletal muscle cells, myogenic cells, we irradiated them with microbeams of protons or α-particles and followed the accumulation of DDR proteins at localized irradiation sites. Thereby, we showed that myoblasts, proliferating myogenic cells, repair local IR-induced DNA damage through both non-homologous end-joining and homologous recombination with different recruitment dynamics depending on the characteristics of ionizing particles (type, energy deposition and time after irradiation), whereas myotubes, post-mitotic myogenic cells, display globally reduced DNA damage response.

## Introduction

DNA is considered as the primary cellular target of ionizing radiation (IR)-induced biological effects (Han & Yu, 2009). IR generates a wide spectrum of DNA lesions, among which DNA double-strand breaks (DSBs) play a key role in the induction of cell cycle arrest and cell death. Indeed, DSBs are the most deleterious DNA lesions for genome integrity and stability (Zhao et al, 2020). In mammalian cells DSBs are repaired through four distinct pathways, primarily by two main mechanisms: classical non-homologous end-joining (c-NHEJ or NHEJ) and homologous recombination (HR). Additionally, residual DSBs, which could not be repaired by NHEJ and HR, can be processed through single-strand annealing (SSA) and alternative end-joining (alt-NHEJ) (Zhao et al, 2020; van de Kamp et al, 2021). While NHEJ is known to be active during the entire cell cycle, HR is limited to S-G2 phases, as it requires template DNA for resynthesis of damaged region guided by homologous sequences on the sister chromatid (Beucher et al, 2009). The principal repair pathway of IR-induced DSBs is reported to be NHEJ (Beucher et al, 2009), which is initiated by the KU70/KU80 heterodimer recruitment and formation of DNA-PK complex with DNA-PKcs (DNA-PK catalytic subunit) (Nick McElhinny et al, 2000; Ahnesorg et al, 2006). Despite this, the radiosensitivity of cells with deficiency in HR-mediated DSB repair (Grosse et al, 2014; Gerelchuluun et al, 2015; Bright et al, 2019) suggests that HR could also be implicated in the repair of DNA damage induced by low or high linear energy transfer (LET) particles.

Nowadays, along with new technologies, new protocols and modalities of radiotherapy are introduced for preclinical studies and clinical use. Depending on the type and/or invasiveness of cancer to be treated, radiotherapy options range from photon (X-ray, γ-ray)- to charged particle (proton, carbon)-treatments. Today, the 5-year survival rate is 70 % and above for certain types of cancer (e.g. prostate cancer, non-Hodgkin lymphoma, kidney and renal pelvis cancer, ovary cancer, breast cancer) (Siegel et al, 2020). Consequently, along with increased survival rate, late onset side effects of cancer therapies are observed (Paulino, 2004; Zhang et al, 2015). Accordingly, skeletal muscle (i.e., myofibers), which makes up to approximately 40 % of the body mass (Frontera & Ochala, 2015), is one of the most liable organs to be exposed to IR as an adjacent healthy tissue during radiotherapy. Despite that skeletal muscle tissue is considered to be radioresistant (Olivé et al, 1995; Coquard, 1997), many studies show that, in long term, muscle surrounding the radiotherapy target displays physiological sequelae depending on the dose, frequency, or quality of IR (Cui et al, 2016; D’Souza et al, 2019). Such muscular problems developed upon radiotherapy include muscle wasting, cachexia, contractures, malfunctioning and weakness, and affect the life quality of patient. Despite their clinical relevance, the molecular and mechanistic origins of radiation-induced muscle aftereffects are still poorly understood.

Skeletal muscle cells originate from unipotent muscle stem cells (Seale et al, 2001). In developing juvenile organisms muscle stem cells are mostly active and proliferative to contribute for skeletal muscle growth (Gattazzo et al, 2020), whereas in adults, they are quiescent and are called satellite cells (SCs), since they are located at the periphery of myofibers between the basal lamina and the sarcolemma. The SCs only activated for self-renewal, muscle homeostasis and regeneration (Collins et al, 2005). Thus, it is only upon activation that SCs proliferate in the form of myoblast, differentiate and fuse into multi-nucleated myotubes and mature into myofibers (Collins et al, 2005). In order to increase our knowledge of late-onset consequences of radiotherapy on muscle, it is therefore important to assess the efficiency of DNA damage repair mechanisms specific to multi-nucleated myofibers as well as its mono-nuclear precursors cells, myoblasts. Previously, it has been reported that adult muscle stem cells have more efficient and accurate DNA repair capacity than their committed progeny (Vahidi Ferdousi et al, 2014). Recently, we have also shown that multi-nucleated skeletal myofibers, generated from muscle stem cells of neonatal mice or immortalized adult myoblasts, have a reduced DNA damage response (DDR) in comparison to their precursors, myoblasts upon x-ray or α-particle irradiation (Sutcu et al, 2024). Additionally, our previous findings suggest that repair of DSBs induced either by x-ray or α-particle irradiation in mono and multinuclear myogenic cells is essentially carried out by NHEJ (Sutcu et al, 2024) as in their progenitor, muscle stem cells (Vahidi Ferdousi et al, 2014; Sutcu et al, 2023)

Due to the increasing, though still marginal, use of protons (proton therapy) and alpha emitters (nuclear medicine), and based on our recent findings on the DDR in myogenic cells upon α-particle irradiation, we performed a study of the DDR in myogenic cells according to the type and characteristics of ionizing particles delivered and the energy deposited. Therefore, taking advantage of ion microbeam technology (Vianna et al, 2022), we delivered different numbers of 4 MeV protons and 6 MeV α-particles to induce local DNA damage at the sub-nuclear level within nuclei of myogenic cells and we followed the recruitment kinetics of some DDR proteins coupled to GFP throughout myogenic differentiation. Thereby, we found that myoblasts orchestrate IR-induced DSB repair with dynamics that depend on the type of high LET particle used, proton or α-particle, while differentiated myotubes remained with declined DNA damage response as was previously reported (Sutcu et al, 2024).

## Results

### The dynamic behavior of KU80-GFP at the DNA damage site depends on the number of particles delivered

To assess initial IR-induced DNA damage response of myogenic cells, we used the immortalized murine C2C7 cells stably expressing GFP-tagged proteins involved in DDR: KU80-GFP, 53BP1-GFP and HP1α-GFP (Sutcu et al, 2024). Immunoblot analyses of whole-cell extracts showed that all GFP-tagged proteins used in this study were expressed at least at a one-to-one ratio, or even less (Fig. S1, A) in stably transfected myoblasts and their committed progeny, myotubes (Fig. S1, B). In addition, no free GFP molecules were detected, while a very weak quantity of truncated products for 53BP1-GFP was found in transfected myogenic cells (Fig. S1, A, comparison between upper and lower panels for each protein). Moreover, we confirmed the nuclear localization of all GFP-tagged constructs used in this study (Fig. S2, A-C), as previously described in literature (Minc et al, 1999; Matsuura, 2019).

To perform *in situ* localized irradiations within the nuclei of living cells, we used the microbeam MIRCOM facility (Bobyk et al, 2022; Sutcu et al, 2024). This microbeam delivers a controlled number of a given particle. Here, we targeted by either 1,000; 200 or 50 α-particles of 6 MeV, or 10,000; 2,500 and 500 protons of 4 MeV in order to have comparable deposited energy between these two types of particles at the localized irradiation sites (Materials and methods and Table S1). To investigate the dynamic of NHEJ upon localized α-particle and proton irradiation, we first monitored the accumulation of KU80-GFP protein at the locally irradiated zones over a time frame of 5 min in myoblasts (Fig. 1, A-B) and myotubes (Fig. S3, A-B). We found that in KU80-GFP-expressing C2C7 cells at the myoblast state, the recruitment of KU80-GFP to DNA damage sites shows similar kinetics upon localized irradiation with 1,000 α-particles (Fig. 1, C) or 10,000 protons (Fig. 1, D), leading to the same deposited energy (Table S1) within the targeted zone. Indeed, immediately after irradiation by 1,000 α-particles or 10,000 protons, KU80-GFP accumulated at the sites of DNA damage, reaching a maximum level of recruitment within 60 s followed by its gradual release (Fig. 1, C-D).

**Figure 1.**
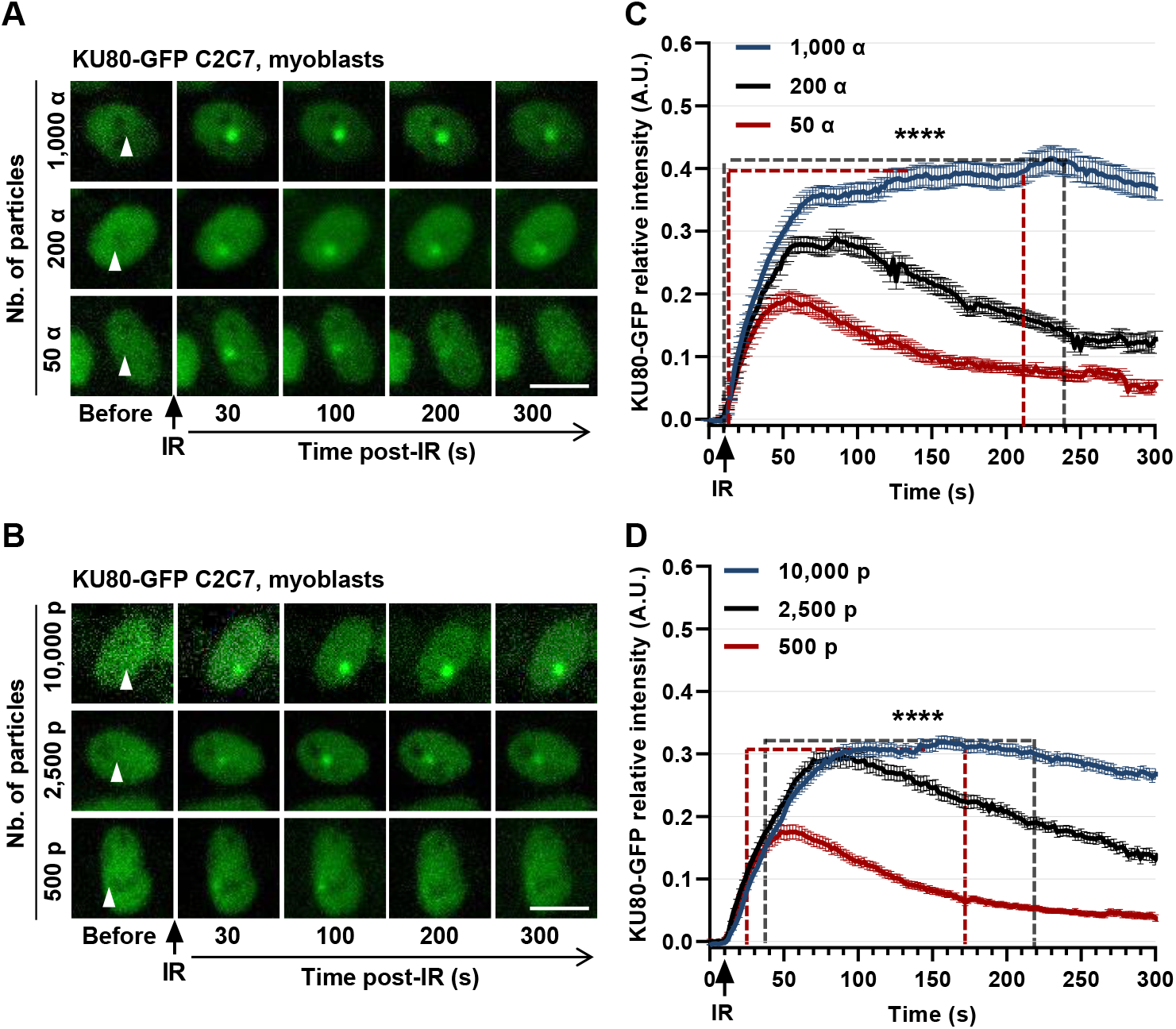
Recruitment kinetics of NHEJ factor KU80-GFP to the site of localized α-particle and proton irradiation within nuclei of proliferating myoblasts. **A, B** Representative images of KU80-GFP recruitment in stably KU80-GFP-expressing C2C7 myoblasts upon localized irradiation (white arrowhead) with the indicated number of α-particles (A) and protons (B). The irradiation was applied at t = 10 s. Scale bar represents 10µm. **C, D** Kinetic curves of KU80-GFP recruitment obtained from A and B upon local irradiation with reducing number of α-particles (C) and protons (D) delivered. Error bars represent the SEM obtained from N= 3 independent experiments with 65 - 210 nuclei / condition analyzed in C and 42 - 175 nuclei / condition analyzed in D. Statistical analysis was performed by using generalized additive models (GAM) framework model and comparing each time point. Dashed boxes represent the region of slopes with significant difference between the kinetic curves. Color code of boxes represents the same color kinetic curve vs curve of highest number of particle irradiation. **** p< 0.00001.

In contrast, in KU80-GFP-expressing C2C7 cells at the myotube state, we found a reduced recruitment of KU80-GFP at the sites of DNA damage induced by a 1,000 α-particles irradiation in comparison to their progenitors, myoblasts (Fig. 1, A and C vs Fig. S3, A and C), as we previously reported in (Sutcu et al, 2024). Likewise, we observed that this was also the case upon 10,000 protons irradiation (Fig. 1, B and D vs Fig. S3, B and D), albeit to a lesser extent. We verified that this highly decreased recruitment of KU80-GFP at localized irradiation sites was not due to a lower amount of this protein in myotubes compared to myoblasts (Fig. S1, A and C,and Fig. S2, A and D). Indeed, the measure of the steady-state levels of KU80-GFP in stably transfected myoblasts and their committed progeny, myotubes, by direct detection of GFP signal (Fig. S2, A and D), showed comparable levels of KU80-GFP proteins in myotubes compared to myoblasts. These data demonstrate that the highly decreased accumulation of KU80-GFP at localized irradiation sites observed in myotubes was not because of a limited amount of this protein.

Next, to evaluate whether the KU80 recruitment to irradiation-damaged DNA is function of the level of deposited energy by ionizing particles, we reduced the number of particles delivered (i.e., 200 and 50 α-particles or 2,500 and 500 protons). In KU80-GFP-expressing C2C7 myoblasts, we thus observed a decrease in the accumulation of KU80-GFP at localized irradiation sites (Fig 1, C-D), transiently reaching to a plateau earlier, followed by its almost immediate release in comparison to irradiations with highest number of particles (i.e., 1,000 α-particles and 10,000 protons). Furthermore, the differences in KU80-GFP recruitment kinetics observed as a function of the number of delivered particles, although modest, are statistically significant, particularly upon α-particle irradiation (Fig. 1 C-D). And as above, KU80-GFP recruitment in myotubes still remains weaker than in myoblasts even upon lower number of α-particles (Fig. S3, A and C) or protons (Fig. S3, B and D) delivered. In myotubes, despite lower KU80-GFP accumulation than in myoblasts, whatever the nature and number of particles delivered, KU80-GFP is retained for longer at localized irradiation sites (Fig. S3, C and D), and especially upon targeted proton irradiation (Fig. S3, D).

Together, our results show for the first time that in myogenic cells the accumulation and disappearance of KU80-GFP proteins at localized irradiation sites are correlated with the number of particles delivered (Fig. 1, C-D), and consequently with the deposited energy and thus with the number and complexity of DSBs induced, as estimated by the Geant4-DNA simulation tool named “dsbandrepair” (Table S1). Indeed, in myoblasts, the faster disappearance of KU80-GFP upon low number of particle irradiation compared to the one with high number of particles seem to correlate with amount of induced DNA damage (Table S1). On the contrary, in myotubes, reduced but persisting KU80-GFP at irradiation-induced DNA damage sites (Fig. S3), which was previously reported to be associated with high turn-over of KU80 proteins (Sutcu et al, 2024), potentially suggests an insufficient amount of KU80 loaded and kept on damaged DNA to process DNA lesions. Additionally, in the absence of other repair pathways, notably HR and reduced BER, as previously demonstrated in (Sutcu et al, 2024), myotubes may exhibit globally weaker DDR compared to myoblasts, possibly impairing and delaying KU80-dependent DSBR.

### The accumulation rate of 53BP1-GFP at DNA damage sites is inversely proportional to the number of particles delivered

53BP1 is a key protein mediating the signaling of DSBs, and orchestrating the choice of DSBR pathways (Bunting et al, 2010) by limiting DNA end-resection and favoring NHEJ (Callen et al, 2020). To assess the initial response of 53BP1 to irradiation, we induced local DNA damage by microbeam irradiation within stably 53BP1-GFP-expressing C2C7 cells and monitored its recruitment to micro-irradiation sites in the time frame of 5 min (Fig. 2, A-B).

**Figure 2.**
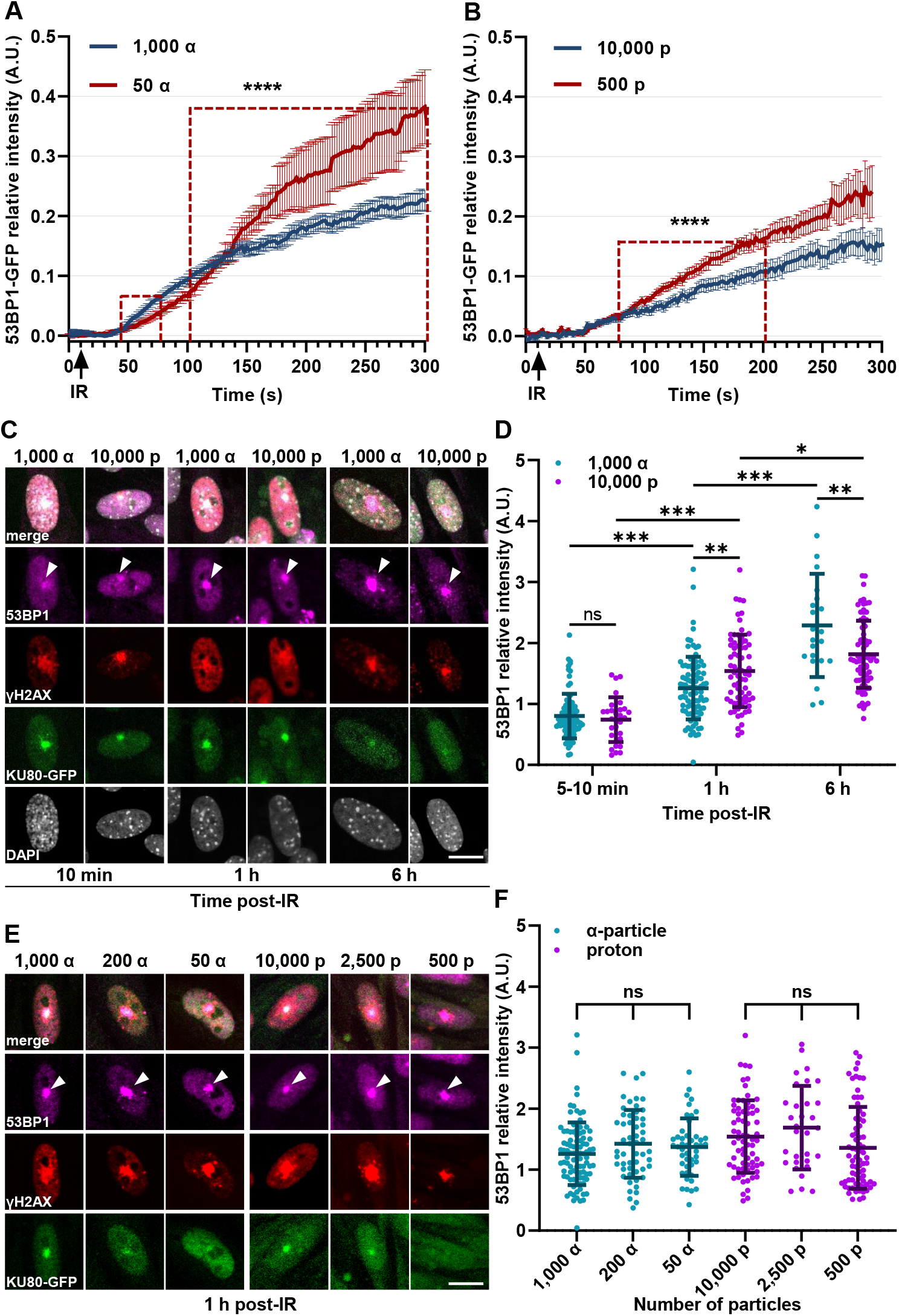
Response of DDR protein 53BP1 upon localized α-particle and proton irradiation within nuclei of myoblasts. **A, B** Kinetic curves of 53BP1-GFP recruitment in stably 53BP1-GFP-expressing C2C7 myoblasts upon 1,000 and 50 α-particles irradiation (A), or upon 10,000 and 500 protons irradiation (B). N= 3 independent experiments, mean ± SEM from 71 – 109 nuclei / condition analyzed in (A) and 23 - 32 nuclei / condition analyzed in (B). Statistical analysis performed by using GAM framework model and comparing each time point. Dashed boxes represent the region of slopes with significant difference between the kinetic curves. Color code of boxes represents the same color kinetic curve vs curve of highest number of particle irradiation. **** p< 0.00001. **C, E** Representative images of stably KU80-GFP-expressing proliferative C2C7 myoblasts immunostained with antibodies against 53BP1 and the phosphorylated form of the histone variant H2AX (γH2AX) at the indicated time-points after localized irradiation (white arrowhead) by 1,000 α-particles and 10,000 protons (C), and 1h after localized irradiation (white arrowhead) by 1,000; 200 and 50 α-particles or by 10,000; 2,500 and 500 protons (E). DNA is counterstained with DAPI. Scale bar, 10 µm. **D, F** Relative fluorescence intensities of 53BP1 staining at local irradiation sites within nuclei of cells from (C) shown in (D) and from (E) shown in (F). N= 3 independent experiments with 22 - 94 nuclei / condition analyzed in (D) and 31 - 94 nuclei / condition analyzed in (F). Data are presented in scatter dot-plots, representing the relative 53BP1-GFP fluorescence intensities at irradiation sites. Bold bars represent the mean and error bars the standard error of the mean (SEM). Statistical significance is obtained by 1-way ANOVA with post-hoc Tukey’s multiple comparison test. p-value are ns p> 0.05, * p< 0.05, ** p< 0.01, *** p< 0.001.

In myotubes we demonstrated that there is almost no initial response of 53BP1 upon 10,000 protons irradiation (Fig. S4, A and C), strengthening our previous data (Sutcu et al, 2024), which have showed this near absence of 53BP1 recruitment upon 1,000 α-particles irradiation. We confirmed that this is not due to a limiting amount of 53BP1-GFP in myotubes (Fig. S1, A and C, and Fig. S2, B and E). However, surprisingly, by reducing the α-particles triggered to 50 we observed a weak but statistically significant increase in the initial 53BP1-GFP recruitment to local IR-induced DNA damage sites in myotubes (Fig. S4, A). Even more remarkably, in myoblasts we also found that lowering the number of particles to 50 α-particles or to 500 protons induces within 5 min an initial recruitment of 53BP1-GFP to irradiation sites almost 2-fold higher than that induced by 1,000 α-particles or 10,000 protons (Fig. 2, A-B). Additionally, unlike KU80-GFP, 53BP1-GFP in myoblasts continues to accumulate and does not reach a plateau in this time frame of 5 min (Fig. 2 vs Fig. 1).

Therefore, to follow 53BP1 recruitment beyond the first 5 - 10 min to IR-induced DNA damage sites and to compare its kinetics to that of KU80, we immuno-labelled KU80-GFP-expressing cells with antibodies against 53BP1 and the phosphorylated form of the histone variant H2AX (known as γH2AX), used here as a DSB biomarker (Goodarzi & Jeggo, 2012). We thus analyzed the fluorescence intensity of their immunostaining at localized sites of irradiation. In KU80-GFP-expressing myoblasts locally irradiated with 1,000 α-particles or 10,000 protons we confirmed that 53BP1 recruitment continues to increase at IR-induced DNA damage sites up to 6 h post-irradiation (Fig. 2, C-D), while KU80-GFP and γH2AX fluorescent signals gradually and simultaneously disappear (Fig. 2, C). In addition, 1h post-irradiation we found that myoblasts irradiated by 10,000 protons have a higher signal of 53BP1 at irradiation sites than those irradiated by 1,000 α-particles (Fig. 2, D). Unexpectedly, the intensity of 53BP1 signal at irradiation sites 1 h post-irradiation is the same regardless of the number of particles targeted (Fig. 2, E-F, α-particles: 1,000 vs 200 and 50; and protons: 10,000 vs 2,500 and 500). In contrast, KU80-GFP and γH2AX signals diminish proportionally with the reduction in the number of particles used for irradiation (Fig. 2, E), consistent with our KU80-GFP recruitment kinetics described above (Fig. 1). At 6 h post-irradiation myoblasts irradiated with 1,000 α-particles show significantly higher 53BP1 intensity at irradiation sites than those irradiated by 10,000 protons (Fig. 2, C-D). This difference indicates that 53BP1 is released faster from DNA following irradiations with protons than irradiations with α-particles, for a same deposited energy at the localized irradiation site.

In contrast, in myotubes the 53BP1 recruitment to irradiated sites is still delayed and strongly reduced even up to 6 h post-irradiation (Fig. S4, B-C). These data indicate that 53BP1 recruitment to IR-induced DNA lesions declines along the myogenic differentiation, thus reinforcing our previous published data (Sutcu et al, 2024).

Taken together, our results show in myoblasts that a low particle number induces faster 53BP1 recruitment to localized irradiation sites than high particle number during the early DDR stages (i.e, within the first 5 min upon irradiation). Then, at 1 h post-irradiation the accumulation of 53BP1 at damaged DNA site is similar, regardless of the number of particles used. Finally, in myoblasts the release of 53BP1 from irradiation-damaged DNA is faster upon localized irradiation with protons than upon the one with α-particles, when the highest numbers of particles are delivered (1,000 α-particles vs 10,000 protons). This dynamic behavior at late stages of DSBR can be linked to a higher yield of DSBs (Table S1) induced by α-particle micro-irradiation than by proton micro-irradiation.

### NHEJ and HR balance in the processing of ionizing particle-induced DSBs in myoblasts

Our above-mentioned results show that within 5 - 10 min post-irradiation the 53BP1 recruitment to damaged DNA sites is inversely correlated with the number of particles delivered (Fig. 2, A-B). As 53BP1 plays a major role in the choice of DSBR pathways (Callen et al, 2020), we next investigated the dynamic behavior of both NHEJ and HR key factors upon localized charged particle irradiation only in myoblasts, as myotubes have no HR activity (Sutcu et al, 2024). This absence of HR activity in myotubes was also confirmed by immunoblotting with almost no detection of RAD51 protein (Fig. S1, B), which is central and pivotal HR factor (Sung & Robberson, 1995; Baumann & West, 1998). Therefore, we analyzed in KU80-GFP-expressing C2C7 myoblasts the local enrichment of γH2AX as DSB marker (Goodarzi & Jeggo, 2012) together with the recruitment of KU80-GFP as NHEJ actor, and RAD51 as HR factor, by direct GFP detection and indirect immunofluorescence at different time-points after targeted irradiation with protons or α-particles (Fig. 3).

**Figure 3.**
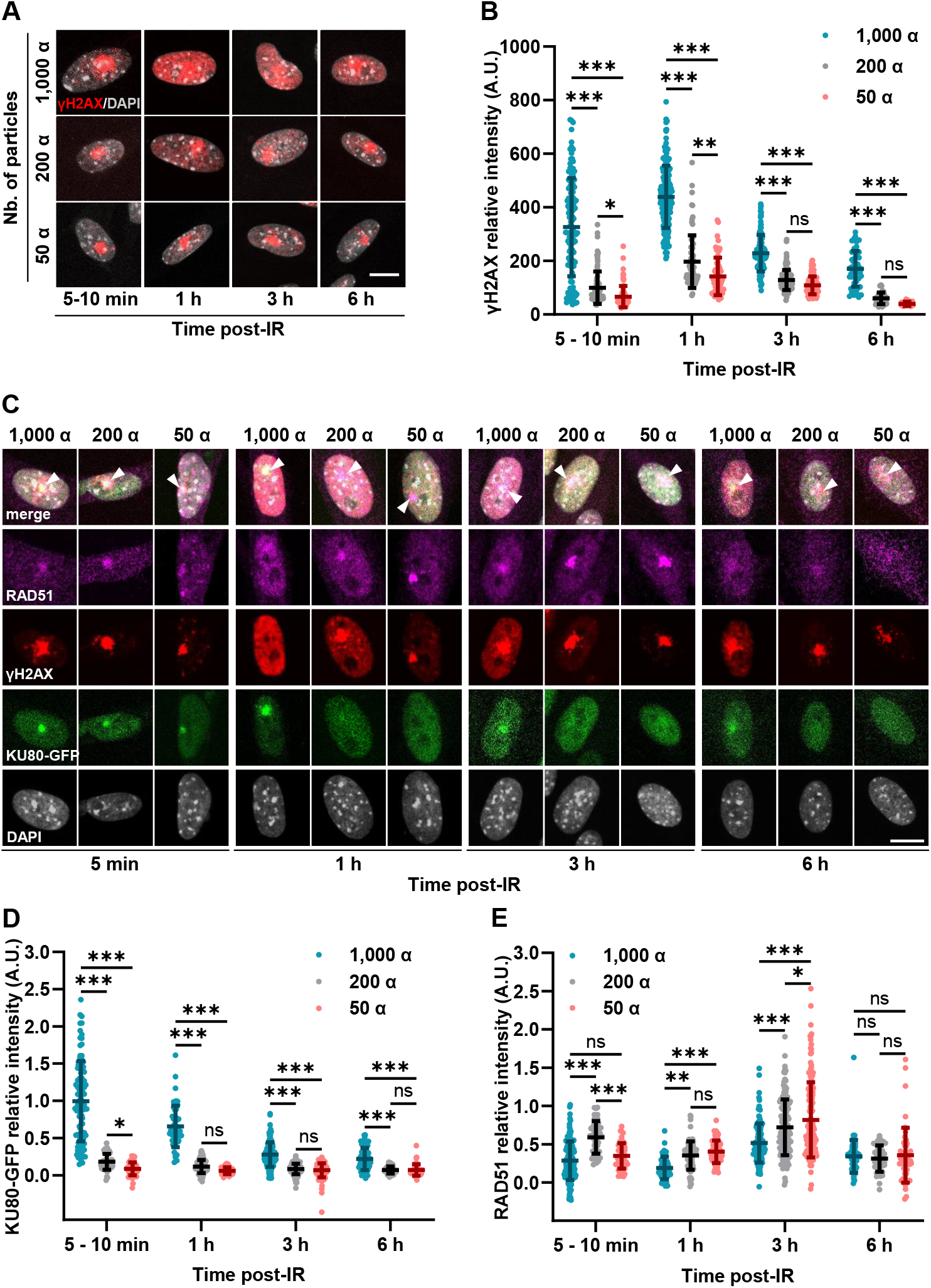
DSB repair mechanisms taking role in myoblasts upon α-particle micro-irradiation with reducing number of particles. **A** Representative images of nuclear γH2AX pattern at the indicated time-points upon micro-irradiation with 1,000, 200 and 50 α-particles in stably KU80-GFP-expressing C2C7 myoblasts immunolabelled against γH2AX. DNA counterstained with DAPI. Scale bar, 10 µm. **B** Mean relative nuclear fluorescence intensities of γH2AX within nuclei of myoblasts from (A) at indicated time-points upon micro-irradiation with the indicated α-particle number. N= 3 independent experiments with 50 - 177 cells / condition analyzed. **C** Representative images of stably KU80-GFP-expressing myoblasts immunolabelled against the HR factor RAD51 and γH2AX at the indicated time-points upon irradiation (white arrowhead) with the indicated number of α-particles. DNA counterstained with DAPI. Scale bar, 10 µm. **D, E** Mean fluorescence intensities of recruited KU80-GFP (D), and immunostaining with anti-RAD51 antibody (E) at the indicated time-points after micro-irradiation with the indicated number of α-particles. N= 3 independent experiments, with 51 - 164 cells / condition analyzed. In scatter dot-plots (B, D, E), are represented the relative fluorescence intensities of nuclear γH2AX staining (B) and, KU80-GFP (D) and RAD51 staining (E) at irradiation sites. Bold bars represent the mean, error bars the SEM. Significance by 2-way ANOVA with post-hoc Tukey’s multiple comparison test against the number of α-particles at each indicated time-points. The p-value are ns p> 0.05, * p< 0.05, ** p< 0.01, *** p< 0.001.

In myoblasts we first observed a pan-nuclear γH2AX staining appearing 1 h after localized irradiation with 1,000 α-particles and being resolved at later time-points (Fig. 3, A). These results are consistent with previous published studies using charged particle irradiation (Meyer et al, 2013; Horn et al, 2015). Furthermore, this pan nuclear signal decreases with the number of delivered particles and is visible as soon as 200 α-particles are delivered (Fig. 3, A). Despite that, to avoid introducing a bias in the quantification of fluorescence signal intensity, we analyzed the whole nuclear γH2AX signal at different time-points after localized irradiation. Accordingly, at each time-point we clearly observed a reduction in the γH2AX signal as a function of deposited energy (Fig. 3, B), directly correlating with a decrease in the number of particles delivered, from 1,000 to 50 α-particles, and with the time elapsed after micro-irradiation, from 1 to 6 h, suggesting an ongoing DSBR. To confirm the correlation of γH2AX fluorescence intensity with the amount of DNA damage induced by a number of particles targeted, we also assessed the total nuclear γH2AX signal in the C2C7 cells irradiated with intermediate number of α-particles (i.e., 500 and 100 α-particles; Fig. S5, A). One-hour post-irradiation, we found that the γH2AX signal is indeed the highest after 1,000 α-particles irradiation and it significantly and continuously decreases in a correlated manner with the reduction of the number of particles delivered, from 500 to 50 α-particles (Fig. S5, A).

Within the first 10 minutes post-irradiation with 1,000 α-particles, the KU80-GFP signal measured at localized irradiation sites is 5- and 10-fold higher compared to that observed following micro-irradiation with 200 and 50 α-particles, respectively (Fig. 3, C-D). Afterwards, the intensity of KU80-GFP signal (Fig. 3, C-D) decreases in a similar manner to that of γH2AX (Fig. 3, B-C). Six hours post-irradiation with 1,000 α-particles, we observed a clear decrease in KU80-GFP signal at localized DNA damage site. However, the fluorescence signal intensity detected after irradiation with lower number of particles (i.e., 200 and 50 α-particles) at the irradiation site was too weak (under our technical imaging conditions) to distinguish any significant difference from 1 h post-irradiation onwards (Fig. 3, C-D and Fig. S5, B). Contrary to KU80-GFP, following irradiation with 1,000 α-particles the RAD51 fluorescence intensity at the localized irradiation site remains relatively low until 1 h post-irradiation (Fig. 3, C-E). Surprisingly, decreasing the number of α-particles from 1,000 to 50 led to an increase in the RAD51 signal intensity at localized irradiation sites, inversely correlating with the KU80-GFP signal and the number of particles delivered (Fig. S5, B).

Three hours post-irradiation, for all of the irradiations with α-particles (1,000; 200 or 50), the RAD51 signal reaches a maximum, with a fluorescence level 2 folds higher, which then is lower 6 h post-irradiation (Fig. 3, C and E). Additionally, in order to better characterize the correlations between the KU80-GFP, γH2AX and RAD51 intensity levels, the number of α-particles targeted and the time after irradiation described in Figures 3 and S5, we performed partial least squares-discriminant analysis (PLS-DA) (Fig. S6, A-B) by taking advantage of having the measured fluorescence intensities of RAD51 and KU80-GFP at the irradiation sites and of nuclear γH2AX for each individual cell. With these cellular identities, PLS-DA adequately classified the cell groups depending on the active DSBR mechanisms (RAD51 and KU80-GFP recruitments) and dynamics (γH2AX) upon irradiation with different number of α-particles (Fig. S6, A, C and E) and throughout time post-irradiation (Fig. S6, B, D and E). Altogether, our data confirmed that in myoblasts NHEJ is the principal pathway involved in the repair of DSBs induced by α-particle irradiation. However, HR plays an important role at the late DSBR stages, in agreement with a potential switch of DSBR pathways from NHEJ to HR as described in (Shao et al, 2012). Surprisingly, upon local irradiations with low number of α-particles, the implication of HR seems to be sooner than for irradiation with 1,000 α-particles.

As DSBR mechanisms depend on the cell cycle (Kakarougkas & Jeggo, 2014), we checked the cell cycle distribution of myoblasts based on their DNA content, by measuring the integrated intensity of DAPI (Toledo et al, 2013; Gruel et al, 2016) within the cell nuclei (Fig. S5, C-D). The distribution of myoblasts in the different phases of the cell cycle at the moment of irradiation was approximately 56.87 ± 5.5% of cells in G1, 3.24 ± 1.5% in S and 13.6 ± 8.7% in G2 (Fig. S5, C). This initial distribution among the cell cycle phases gives the probability of inducing DNA lesions in G1, S, or G2 in myoblasts randomly chosen within the cell monolayer. Within 5 - 10 min after irradiation with α-particles (Fig. S5, Di), the irradiated myoblasts show similar cell cycle distribution as neighboring non-irradiated myoblasts (Fig. S5, C – 5 - 10 min), 62.3 ± 10.7% of cells in G1, 13,3 ± 10.7% in S and 17.3 ± 10.4% in G2 (Fig. S5, Di – 5 - 10 min). Thus, although no obvious change in the cell cycle distribution of irradiated myoblasts is observed within the first 10 min post-irradiation (Fig. S5, D), the targeted cells display strong γH2AX and KU80-GFP signals (Fig. 3 and S5, A-B). Moreover, despite the cell cycle distributions remaining comparable at 1h and 3h post-irradiation, with 33.7 ± 5.05% and 35.5 ± 8.6% of cells, respectively, in S/G2 phases (Fig. S5, Di), when HR-mediated DSBR is active, the mean fluorescence intensity of RAD51 staining at localized irradiation sites increases only at 3 h post-irradiation (Fig. 3, E), compared to that detected 1 h post-irradiation. Together, the cell cycle distribution and PLS-DA analyses suggest that following α-particle irradiation, NHEJ is the initial responding DSBR mechanism, while HR-mediated repair is engaged later, with slower kinetics, which the contribution of the cell cycle progression alone is not sufficient to fully explain.

As differences in DSBR mechanisms and dynamics are observed by reducing energy deposition of α-particles, we next investigated and compared the DDR of myoblasts to DNA lesions of different complexity induced by protons. At localized irradiation sites we analyzed the fluorescence intensities of RAD51 and KU80-GFP as well as the nuclear γH2AX signal from 5 - 10 min to 6 h post-irradiation with 10,000 protons and 1,000 α-particles (Fig. 4, A-C), leading to similar deposited energies (Table S1). Upon localized proton irradiation of myoblasts, RAD51 fluorescence intensity reaches its maximum at 1 h post-irradiation, sooner than upon micro-irradiation with α-particles (Fig. 4, A). Moreover, the decrease in RAD51 signal at localized irradiation sites is faster upon proton irradiation, starting from the time-point 3 h post-irradiation (Fig. 4, A). However, the KU80-GFP recruitment and release kinetics at local DNA damage sites are similar to those observed in myoblasts irradiated with α-particles, but on a smaller scale (Fig. 4, B). Altogether, the faster release of RAD51 and the lower accumulation of KU80-GFP at the local DNA damage site upon micro-irradiation with 10,000 protons in comparison to micro-irradiation with 1,000 α-particles (Fig.4, A-B), together with the more rapid disappearance of γH2AX signal over time after IR (Fig. 4, C), suggest that DNA lesions induced by proton micro-irradiation are repaired faster. Accordingly, the projection of the data obtained in Figure 4 to the previous PLS-DA analysis (Fig. S6) showed distinct scatterplot projections of myoblasts micro-irradiated by protons or α-particles both for the 1 h and 3 h time-points due to strong differences of both KU80-GFP and RAD51 fluorescence identities of cell groups (Fig. S7). The results obtained from the PLS-DA analyses reinforce the idea that the dynamics of DNA repair mechanisms are different depending on the nature of the particle used for irradiation, but also on the number of particles delivered, despite similar cell cycle distribution of α-particle (Fig. S5, Di) and proton (Fig. S5 Dii) irradiated cells.

**Figure 4.**
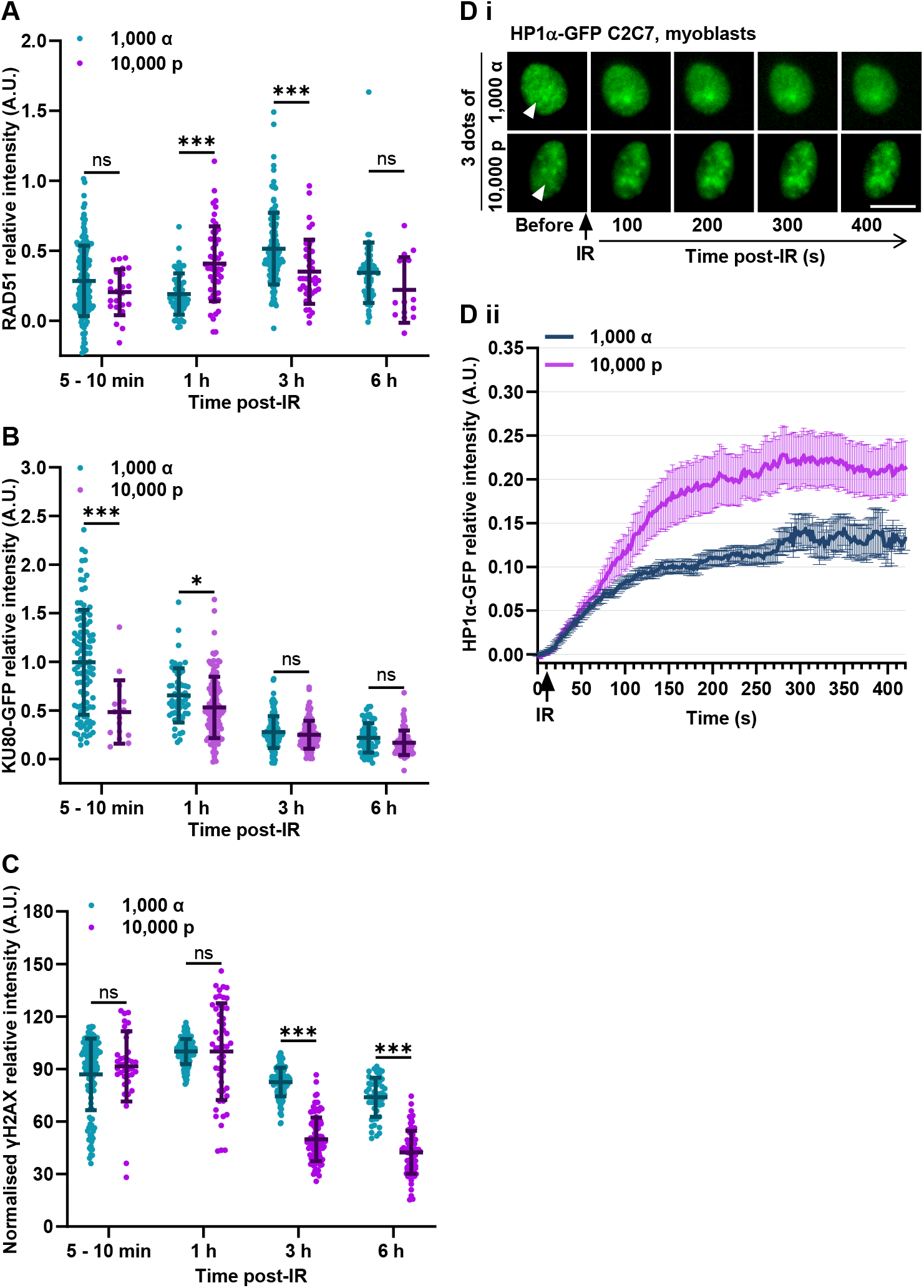
DSB repair upon α-particle and proton irradiation. **A - C** Mean fluorescence intensities of immunolabelled recruited RAD51 (A), recruited KU80-GFP (B) and immunolabelled total nuclear γH2AX (C) at indicated time-points upon α-particle and proton irradiation. N= 3 independent experiments, with 55 – 177 cells / condition analyzed. Data are presented in scatter dot-plots. Bold bars represent the mean, error bars the SEM. Significance by 2-way ANOVA with post-hoc Sidak’s comparison test against the number of particles delivered at each indicated time-points. The p-value are ns p> 0.05, * p< 0.05, ** p< 0.01, *** p< 0.001. **D** Representative images (Di) and kinetic curves (Dii) of HP1α-GFP recruitment in C2C7 myoblasts upon 1,000 α-particles and 10,000 protons micro-irradiation. N= 3 independent experiments, mean ± SEM from 30 - 39 cells / condition analyzed.

Since HR seems to be involved sooner in the process of proton-induced DSBR than that induced by α-particles, we analyzed the behavior of the heterochromatin protein 1 isoform, HP1α. The recruitment of HP1α by p150CAF-1, the largest subunit of histone chaperone chromatin assembly factor 1 (CAF-1), to the laser micro-irradiation induced DNA damage site promotes HR-mediated DSBR (Baldeyron et al, 2011). Thus, to further characterize the cellular response to localized irradiation by charged particles, we analyzed the recruitment kinetics of HP1α within stably HP1α-GFP-expressing C2C7 myoblasts upon irradiation with 10,000 protons or 1,000 α-particles (Fig. 4, Di). In agreement with our data above (Fig. 4, A), HP1α-GFP recruitment to localized irradiation sites is also higher upon proton irradiation than upon α-particle irradiation (Fig. 4, Dii). These results suggest that HR-mediated DSBR takes place earlier upon proton micro-irradiation when compared to α-particle micro-irradiation.

Here we showed that in myoblasts NHEJ is the initial and principal repair pathway to process DNA DSBs induced by localized α-particles irradiation and then there is a switch of repair machinery to HR around 3 h post-irradiation. Additionally, decreasing the α-particle number delivered leads to an earlier involvement of HR in the repair of IR-induced DNA lesions. Upon proton irradiation, unlike α-particle-induced DNA damage, HR seems to take role in the DNA damage processing much earlier, 1 h post-irradiation, and at a faster pace since a decrease in RAD51 staining is already observed at 3 h post-irradiation.

## Discussion

Along with increasing cancer survival rates, the late side effects of radiotherapy treatments on muscle tissue are becoming more apparent (Paulino, 2004; Zhang et al, 2015; Siegel et al, 2020). Muscle is one of cancer-adjacent healthy tissue majorly exposed during IR treatment leading to impaired life quality of patients. This is reported to be more severe in pediatric patients who still have an ongoing muscle development (Goodenough et al, 2021), and thus their muscle tissue that includes all forms of myogenic cells: quiescent, active muscle stem cells; cycling and differentiating precursor cells, myoblasts; and post-mitotic myofibers, as was reported in mice (Gattazzo et al, 2020). Previously we have shown that myotubes have reduced DDR upon irradiation and slower DSB repair capacity in comparison to their precursor cells, myoblasts (Sutcu et al, 2024). However, the repair mechanisms acting in proliferating myoblasts remained poorly characterized. Therefore, we investigated the DDR in myogenic cells upon irradiation with different types of charged particles by using ion microbeam technology (Vianna et al, 2022), offering the advantage of delivering controlled number of particles within a sub-nuclear region of cell nucleus. Thereby, we were able to monitor the cellular response to locally induced DNA damage, particularly by ionizing radiation, while most of the published local DNA damage studies are performed by laser micro-irradiation.

### Dose-dependent retention of KU80 at DNA damage site

We found here that the level of KU80 recruited and retained at the local damage site directly correlates with the number of particles delivered within the nuclei of myoblasts. Since the γH2AX signal serves as a marker of the presence of DSBs and its gradual disappearance reflects the kinetics of DSBR, our analyses of the KU80-GFP accumulation and resolution kinetics, demonstrate for the first time that NHEJ-mediated DSB repair can cope with increasing amount of IR-induced DNA lesions.

Additionally, in all of the irradiation conditions, myotubes show a lower recruitment of KU80 to localized irradiation site in comparison to myoblasts, strengthening the data of our previous work (Sutcu et al, 2024), but a longer retention of KU80 on damaged DNA. One reason for the longer persistence of KU80 at local DNA damage sites in myotubes could be due to the absence of HR repair pathway, as reported in (Vahidi Ferdousi et al, 2014; Sutcu et al, 2024), leaving myotubes entirely dependent on NHEJ to repair IR-induced DSBs, whereas cycling myoblasts still have the capability of deploying HR.

Finally, as previously reported, the recruitment of KU80 occurs in all myogenic cells and its release from DNA may happen when HR takes place to mediate repair of more complex DSBs (Shibata et al, 2011; Shao et al, 2012). Thus, in myoblasts the faster release of KU80 upon irradiation with smaller number of particles could be due to the generation of smaller number of DSBs of lesser complexity, which may explain a faster repair, and a potentially earlier switch of DSBR mechanisms when NHEJ is no longer sufficient to cope with the complexity of remaining IR-induced DNA lesions.

### Recruitment of 53BP1 correlates with the presence of HR activity in myogenic cells

53BP1 is a crucial DDR protein, which has multiple roles in DSBR (Shibata & Jeggo, 2020). It is well established that in proliferating cells 53BP1 recruitment to DSBs promotes DSBR through NHEJ by restricting the 5= DNA end resection and thus preventing the implementation of HR-mediated DSBR (Bunting et al, 2010; Callen et al, 2020). Here we showed that in post-mitotic myotubes 53BP1 recruitment is very low or even almost non-existent at the charged particle irradiation sites. This reduced 53BP1 recruitment in myotubes could be the consequence of a general decline in NHEJ along myogenic differentiation, as shown above with KU80 (Sutcu et al, 2024). However, we can also hypothesize that in absence of functional HR in myotubes (Sutcu et al, 2024), characterized by a very weak expression of RAD51, 53BP1 is not required, since there is no need to counteract HR implementation to promote NHEJ. Indeed, NHEJ can efficiently occur in absence of 53BP1 for most DSBs (Ward et al, 2004). However, besides its role in the choice of DNA repair pathway, 53BP1 recruitment also induces the chromatin relaxation for the ATM-dependent slow-kinetic DSBR at heterochromatin regions (Noon et al, 2010). Interestingly, it was reported that in myotubes the activity of ATM and its downstream effectors is low (Latella et al, 2004), which would leave the DNA lesions in heterochromatin regions unprocessed or difficult to process. Accordingly, our observations suggest that in myotubes the very late recruitment of 53BP1 at 6 h post-irradiation would probably provide a protection against unprocessed broken DNA ends, which was reported to be an ATM-independent mechanism (Rybanska-Spaeder et al, 2013). Likewise, although in myoblasts 53BP1 accumulation at DNA damage sites is already higher than in myotubes within 5 – 10 min after micro-irradiation, it also gradually reaches to its maximum later at 6 h post-irradiation when KU80 becomes undetectable.

Slower recruitment of 53BP1 upon high LET micro-irradiation with high number of charged particles suggests that 53BP1 is potentially dispensable at the first steps of DSBR in myogenic cells but it is required in later stages for the processing of more complex DSBs. We have also noticed that 53BP1 accumulation is higher within 5 – 10 min following irradiation with a smaller number of particles, either proton or α-particle. Concurrently, this was also the case with RAD51, suggesting earlier commitment of HR. Recently, sooner implication of HR was also reported in lymphoblastoid cell lines upon exposure to low doses of X-ray irradiation (Mladenov et al, 2022). Moreover, RAD51 recruitment has also been correlated with 53BP1 foci formation in human lung epithelial carcinoma A549 cells upon low doses of X-ray irradiation (Mladenov et al, 2020). Besides, in the absence of p150CAF-1-dependent HP1α recruitment, HR was found to be suppressed as well as the recruitment of 53BP1 to laser micro-irradiation-induced DNA damage sites (Baldeyron et al, 2011). Hence, in agreement with the literature, the direct correlation of RAD51 and 53BP1 recruitment to locally damaged DNA sites upon irradiation with charged particle could implicate 53BP1 in the DNA broken end protection from extensive resection, limiting alternative DSBR pathways (i.e., single strand annealing, SSA, or alternative non-homologous end-joining, alt-NHEJ) (Bakr et al, 2016), and allowing the HR to take over. Indeed, upon localized irradiation with high particle number, unlike the faster accumulation of KU80 favoring NHEJ, RAD51 accumulation is delayed to later stages, most likely to process remaining unrepaired DNA broken ends by HR. Consolidating these data, the “dsbandrepair” tool we used to estimate that the yield of DSBs indicates that the proportion of complex DSBs increases with the number of ionizing particles targeted (Table S1). Thus, although it is contradictory to its generic role in favoring NHEJ, our data imply a novel role for 53BP1 in broken DNA end protection, if not favoring, at least to maintain the DNA ends to be processed by HR. In agreement with this, the unexpected behavior of 53BP1 in myotubes upon local DNA damage induction states that 53BP1 is dispensable in the implementation of NHEJ-dependent DSBR in myogenic cells.

### HR is implicated sooner in the processing of DNA lesions induced by protons, which are also repaired faster

There are multiple studies pointing out that irradiations with different types of particles (i.e., species, energy and linear energy transfer of particles) induce different kinds of DNA damage (Bertolet et al, 2023) with different levels of complexity (i.e., different types of DNA lesions in the same DNA region) and clustering (i.e., accumulation of DNA damage in the same DNA region) ((Nikjoo et al, 2001; Rezaee & Adhikary, 2024). Computational approaches (Nikjoo et al, 2001; Rezaee & Adhikary, 2024), which are the only dedicated tools, estimated that irradiations with α-particles lead to a small number of DNA damage sites with concentrated clusters of complex DNA lesions compared to irradiations with protons, which give a higher number of DNA damage sites containing less clusters of complex DNA lesions. Here, with the use of the MIRCOM ion microbeam, we induced, whatever the particle delivered, protons or α-particles, DNA lesions within a very localized subnuclear area, thus potentially more clustered. We used a computational method called “dsbandrepair” (Meylan et al, 2017; Tuan Anh et al, 2024) for estimating the deposited energy of particles delivered by MIRCOM and the radiation-induced DNA damage yields at irradiation sites within myoblasts. This Monte Carlo-derived simulation tool allows, along the track of particles within cell nuclei, the estimation of number of (i) strand breaks (SBs), (ii) simple DSBs (sDSBs) and (iii) complex DSBs (cDSBs) defined herein as a single DSB including one or several SBs (Meylan et al, 2017). The “dsbandrepair” tool has estimated that at comparable deposited energy, localized irradiation with α-particles produced twice more DSBs than localized irradiations with protons (Table S1), and the yield of cDSBs is 10 times more with α-particles than with protons (Table S1).

Accordingly, the detection of histone H2AX phosphorylation (γH2AX) localized only at the irradiation sites suggests the formation of localized irradiation-induced DSBs. However, after delivering at least 200 α-particles we observe that H2AX phosphorylation propagates transiently throughout the whole cell nucleus, in addition to a persistent γH2AX signal at the irradiation sites up to 6 h post-irradiation. In agreement with our previous findings (Sutcu et al, 2024) and the literature (Meyer et al, 2013; Horn et al, 2015), this pan-nuclear γH2AX signal is not linked to the induction of apoptosis (Cook et al, 2009) nor to the presence of DSBs, since, like others (Meyer et al, 2013; Horn et al, 2015), we do not find 53BP1 accumulation outside the localized irradiation sites. This pan-nuclear γH2AX is likely due to a stress signal resulting from an increase in indirect DNA lesions induced by the production of reactive species, and/or increase in the complexity and/or clustering of DNA damage generated by charged particle-irradiation (Meyer et al, 2013; Horn et al, 2015; Răileanu et al, 2022; Bertolet et al, 2023). Nevertheless, although this γH2AX signal does not correspond to high LET IR-induced DSBs, it still directly correlates with the number of charged particles delivered during irradiation and its disappearance with the repair of DNA lesions induced by protons or α-particles. Based on our results, we can hypothesize that the pan-nuclear γH2AX signal could serve as a marker of increased complexity and/or clustering of IR-induced DNA lesions, as long as it does not coincide with the induction of apoptosis. Finally, regardless of the type of charged particle, the robust recruitment and the quick release of KU80-GFP at irradiation sites with low number of α-particles or protons suggest that the easy-to-ligate DNA broken-ends are quickly processed by NHEJ. Then, HR processes the remaining complex DNA lesions, with a sooner implication when irradiation is performed with low number of particles.

Furthermore, the faster disappearance of γH2AX signal after proton irradiation, which we have observed to be concomitant with the faster decrease in KU80-GFP and RAD51 signals at targeted irradiation sites suggest that DNA lesions induced by proton is processed faster than the ones induced by α-particle with comparable energy deposition. Our findings agree with simulation studies (Bertolet et al, 2023) and our stimulation data using “dsbandrepair” tool (Table S1), suggesting that α-particles induce more complex and clustered DNA damage than protons at comparable deposited energies. While NHEJ is thought as the principal DNA repair mechanism for DNA damage induced by charged particle-irradiation, the importance of HR was also recently reported upon proton irradiation (Vitti & Parsons, 2019). Besides, our data show an earlier recruitment and disappearance of 53BP1 upon proton irradiation, in addition to a higher recruitment of HR-promoting factor HP1α, in comparison to α-particle irradiation in myoblasts. Thus, we confirm that HR is actively involved in the proton-induced DNA damage processing alongside with NHEJ, while HR takes place later following α-particle irradiation. These results suggest that DNA damage induced by α-particles would require more DNA end-processing by NHEJ before leaving to HR the repair of the remaining DSBs, leading to a slower resolution of the γH2AX signal (Fig. 5). This could also explain the longer persistence of KU80-GFP at the DNA damage site in HR-deprived myotubes upon irradiation with protons than with α-particles, probably to compensate the lack of HR activity. In conclusion, in proliferating myoblasts the DSBR mechanism engaged depends not only on the deposited energy (i.e., increased number of charged particles), but also on the type of the charged particle and consequently on the amount of complex and/or clustered DNA damage induced (Table S1).

**Figure 5.**
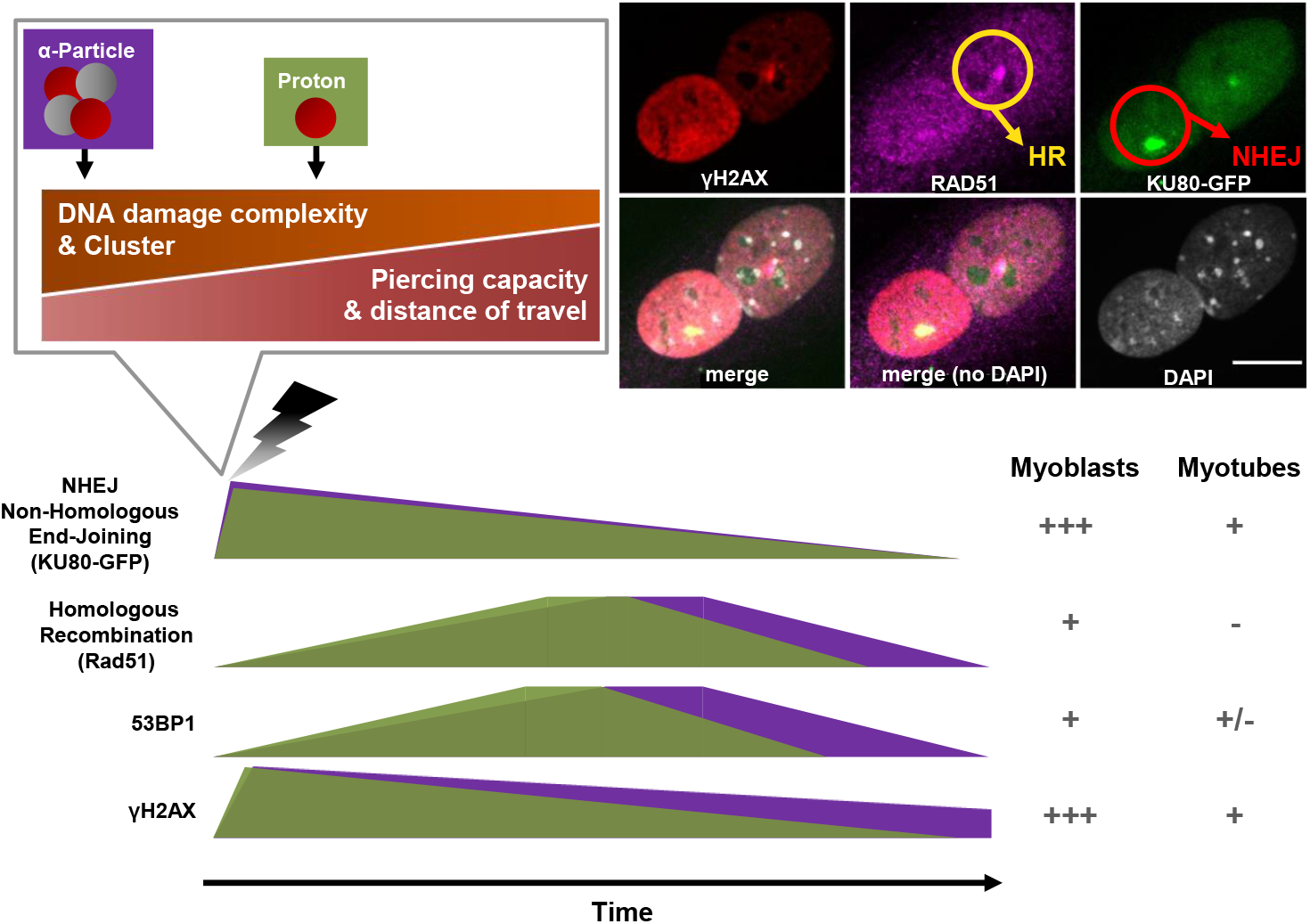
Different DDR dynamics relying on the characteristics of ionizing charged particles in myoblasts. Schematic representation (not in scale) of recruitment and release of NHEJ factor (KU80-GFP), HR factor (RAD51), DDR protein (53BP1) and phosphorylation of histone H2AX (γH2AX) and their degree of implications “+ or −” in myoblasts and myotubes upon α-particle (in purple) and proton (in green) micro-irradiation, hence upon different complexities of induced DNA damage (Bertolet et al, 2023; Rezaee & Adhikary, 2024), against time. P.S. decreasing number of particles delivered also followed similar dynamics as for decreased DNA damage complexity (i.e. proton vs α-particle).

In conclusion, in myoblasts HR is involved later, potentially to repair unprocessed more complex and/or clustered DNA lesions. However, HR also seems to be more strikingly involved in the repair of DNA damage induced by proton irradiation, probably due to the fact that protons in comparison to α-particles induce much less complex DSBs (Table S1). Moreover, myogenic cells seem to repair DSBs with distinct mechanisms that adapt and change throughout myogenic differentiation, in which 53BP1 is dispensable in myotubes and in the early stages of NHEJ-dependent DSBR of myoblasts (Fig. 5).

Finally, while this study provides further understanding on how skeletal muscle cells respond to different radiation qualities that could be used during radiotherapies, it also points out the importance of considering the cancer-adjacent healthy tissue in radiotherapy modalities and treatment planification to be performed on patients. This is even more vital when radiotherapy is particularly combined with chemotherapies, targeting certain DNA damage repair mechanisms and cell cycle control. In that matter, these treatments could also block the recovery of the healthy tissue from induced DNA damage, especially in pediatric patients, who are still physiologically growing and for instance hosting actively cycling myogenic cells apart from differentiated skeletal muscle cells, hence multiple DSBR mechanisms.

## Materials and methods

### Myogenic cell culture

Immortalized myogenic cells, C2C7 were kindly gifted by Miria Ricchetti (Institut Pasteur, Département de Biologie du Développement et Cellules Souches, Laboratoire de Mécanismes Moléculaires du Vieillissement Pathologique et Physiologique, Paris, France). As previously described in (Sutcu et al, 2024), we maintained the C2C7 cells (Pinset et al, 1988) in myoblast state (proliferative state), by culturing in growth medium (GM) based on DMEM Glutamax (Gibco) supplemented with 20 % fetal calf serum (FCS, Eurobio), and 100 U.mL^−1^ penicillin and 100 µg.mL^−1^ streptomycin (i.e., 1 % P/S; Gibco). To generate post-mitotic myotubes, once myoblasts have reached 80 % confluency, we switched the medium to differentiation medium (DM) corresponding to phenol-red free DMEM (Gibco) complemented with 2 % horse serum (HS, Gibco), 1 % P/S, 1 % Glutamax (Gibco), and 1 % Sodium Pyruvate (Gibco). To note, 50 % of DM was refreshed after 3 days and every consecutive day after. We incubated all the cells in a humidified atmosphere at 37 °C with 5 % CO_2_ and 3 % O_2_.

During irradiation experiments on MIRCOM platform, we grew immortalized C2C7 myoblasts and their committed progenies, myotubes on 4 µm thick polypropylene foil (Goodfellow) pre-coated with 10 ng.µL^−1^ of Cell-Tak (Corning) followed by 10 % Matrigel (Corning) in DMEM on special PEEK dishes, as described in (Vianna et al, 2022). On the day of irradiation, we replaced the GM medium with phenol-red free version to reduce potential autofluorescence.

### Plasmids and transfections

The plasmids expressing GFP-tagged protein of interest were kindly provided by Pascale Bertrand (53BP1-GFP; CEA, iRCM/IBFJ, UMRE008 Stabilité Génétique, Cellules Souches et Radiations, Fontenay-aux-Roses, France), Dik C. van Gent (KU80-GFP; Erasmus MC, Departments of Cell Biology and Genetics, Rotterdam, The Netherlands), and G. Almouzni (HP1α-GFP; Institut Curie, UMR3664 Dynamique du Noyau, Paris, France). As previously described in (Sutcu et al, 2024), these plasmids were transfected in C2C7 myoblasts with TurboFect (ThermoFisher Scientific) according to the manufacturer’s instructions. To get successful transfection, C2C7 cells were transfected 1-day post-seeding at about 50 % - 60 % confluency. Then, to obtain stable and homogenous GFP-tagged protein expressing C2C7 lines, 24 h post-transfection the GFP-expressing cells were enriched under 1 mg.mL^−1^ of geneticin (G418 sulfate; Gibco) selection for 10 consecutive days. Then, the GFP-tagged protein expressing C2C7 cells were isolated by FACs sorting and further expanded (Sutcu et al, 2024). To note, GFP-tagged protein-expressing myotubes were obtained by inducing differentiation of their precursors, GFP-tagged protein-expressing myoblasts.

### MIRCOM, Microbeam particle irradiation

We performed the localized irradiation of samples by 4 MeV protons, or 6 MeV α-particles generated from hydrogen and helium ions, respectively, by using the MIRCOM facility, operated by the Institute for Radiological Protection and Nuclear Safety (IRSN) in Cadarache, France (Vianna et al, 2022). This facility is equipped with a focused ion microbeam, which is produced by a 2 MV Tandetron™ accelerator manufactured by High Voltage Engineering Europa B.V. (HVEE, Amersfoort, The Netherlands). It has been designed to perform in-air targeted micro-irradiation of living biological samples with a controlled number of ions and a targeting accuracy of 2.1 ± 0.7 µm with a beam spot diameter of a few micrometers (Vianna et al, 2022).

As protons and α-particles have low penetration capacity and thus short travel distance through matter, the cells are seeded in a specific PEEK cell dish with a 4 µm thick polypropylene foil (Goodfellow) (Vianna et al, 2022). The LET of the particles at the entrance of the samples after going through the extraction window (150 nm), a residual air layer (250 µm), and the polypropylene foil (4 µm) is 84 keV.µm^−1^ for 6 MeV α-particles (Bobyk et al, 2022) and 10 keV.µm^−1^ for 4 MeV protons, calculated with SRIM (Vianna et al, 2022). Thus, by using the computational method called “dsbandrepair” (Table S1), we estimated the specific energy deposited within the targeted volume, and it was approximately 35,04 keV for one 4 MeV proton and 346,2 keV for one 6 MeV α-particle in our experimental conditions of localized irradiation (Table S1). The minimum number of particles targeted is fixed to 50 α-particles and 500 protons, since it is the minimum particle number allowing the detection of RAD51 recruitment under our experimental conditions (data not shown).

During irradiation, myogenic cells are placed under an inverted epifluorescence microscope (AxioObserver™ Z1, Carl Zeiss Microscopy GmbH, Jena, Germany) within a 37 °C heating chamber. The nuclei of GFP-expressing cells are identified and selected for irradiation with a 20x objective (Zeiss LD Plan-NEOFLUAR 20x/0.4 Korr). Then, we targeted the nuclei of selected myogenic cells with the indicated number of protons or α-particles, which was determined by the definition of an opening time of the microbeam on each target, as described previously (Bobyk et al, 2022; Vianna et al, 2022; Sutcu et al, 2024). Thus, the energy deposition of each irradiation with protons or α-particles is directly correlated to the number of particles delivered (Table S1).

To follow the recruitment kinetics of our GFP-tagged proteins of interest, we started time-lapse imaging 10 s before targeted irradiation with the indicated number of particles and recorded images every 2 s with a monochromatic AxioCam™ MRm rev. 3 CCD camera (Carl Zeiss Microscopy GmbH, Germany) using the CRionScan software (Vianna et al, 2022). We recorded images with an exposure time of 800 ms. In total, we kept myogenic cells in the microbeam chamber for less than 30 min.

### Immunostaining and image acquisition

At the indicated time following irradiation, myogenic cells were fixed with 2 % paraformaldehyde (PFA; EMS, Euromedex) in phosphate-buffered saline (PBS; Gibco) for 20 min, and then permeabilized with 0.5 % Triton X-100 (Sigma-Aldrich) in PBS for 5 min. To increase the stringency cells were washed with 0.1% Tween 20 (Sigma-Aldrich) in PBS (PBS-T), then blocked with 5% BSA (Sigma-Aldrich) in PBS-T. Next, myogenic cells were incubated with the indicated primary antibodies overnight at 4°C. After three washes in PBS-T, the cells were incubated with the appropriate fluorophore-conjugated secondary antibodies for 1 h at room temperature (RT). After three washes in PBS-T, myogenic cells were incubated with 0.5 µg/mL DAPI (Molecular Probes) in PBS for 5 min at RT. Finally, after three washes in PBS, myogenic cells on polypropylene foil were mounted between slide and coverslip with Prolong Diamond antifade mounting medium (Molecular Probes). The myogenic cells were imaged with C-Plan Apochromat 63x/1.4 Oil DIC M27 objective under a LSM 780 NLO confocal microscope, piloted with the Zen Black 2011 SP4 software (Carl Zeiss MicroImaging GmbH, Germany).

### Antibodies

Primary antibodies used during immunofluorescence experiments are: rabbit anti-53BP1 (Bio-Techne, Novus Biologicals, NB 100-304; 1:500); mouse anti-γH2AX (Millipore, UpState, 05-636; 1:2,000); rabbit anti-RAD51 (Abcam, ab137323; 1:400).

Secondary antibodies used are: donkey anti-mouse or donkey anti-rabbit coupled to Alexa Fluor 594 or 647 (Invitrogen, ThermoFisher Scientific; 1:1,000). To amplify the GFP fluorescence intensity post-fixation, we used GFP booster nanobodies conjugated with Alexa fluor 488 (Chromotek, gb2AF488; 1:500).

### Quantification and Statistical analyses

All data presented in this paper result from at least three independent experiments.

All the images were processed, analyzed, and quantified by ImageJ software (version 1.53e) (Schneider et al, 2012), and statistical analyses were performed by Prism software (version 10; GraphPad Inc.) and Excel (Microsoft).

As previously described (Sutcu et al, 2024), in order to quantify the fluorescence re-localization of GFP-tagged proteins observed by time-lapse imaging upon MIRCOM irradiation, we manually selected and followed regions of interest (ROI). We measured the mean intensity of ROIs in every image and plotted them against time. To obtain the relative intensity of GFP-tagged repair proteins recruited at the irradiation site, the mean intensity of non-irradiated site was subtracted from ROIs and normalized to mean fluorescence intensity of total nuclei. Then the obtained data were corrected for non-specific fluorescence bleaching and normalized for the fluorescence intensity measured before irradiation.

The smoothing and comparison between the kinetic curves of KU80-GFP and 53BP1-GFP recruitment under different irradiation conditions were performed within the generalized additive models (GAM) framework (Hastie & Tibshirani, 1995) where the smoothing of the KU80-GFP time-kinetic data was performed using penalized splines. The statistical inference on the difference between two kinetic curves was based on the uncertainties of the smoothing process: a difference at a given time-point was considered as significant if its associated confidence interval did not contain zero. This approach permitted us to highlight the significant differences along the time-point values.

Fluorescence intensity analysis over time-points post-fixation was performed in a similar manner to time-lapse image analysis, as described above, on immunolabeled samples. The relative fluorescence intensity of immunolabeled proteins recruited to irradiation sites was obtained by subtracting the mean intensity of non-irradiated sites from the mean intensity of regions of interest (ROIs, irradiated site) - of the same size across all conditions - and normalizing to the mean fluorescence intensity of total nuclei. As a marker of DSB, the appearance and disappearance of the phosphorylation of histone variant H2AX upon induced DNA damage were analyzed over time. Specifically, we measured the mean intensity of immunolabeled γH2AX in the entire irradiated nucleus, minus background intensity.

## Author Contributions

**HHS:** Conceptualization, Data curation, Formal Analysis, Investigation, Methodology, Project administration, Validation, Visualization, Writing – original draft. **ATJ** and **DD:** Investigation, Methodology. **MCM:** Methodology (α-particle and proton irradiation experiments). **FV:** Methodology, Supervision (α-particle and proton irradiation experiments). **YP** and **MAB:** Formal Analysis. **MB:** Supervision, Validation. **CB:** Formal analysis, Funding acquisition, Project administration, Supervision, Validation, Visualization, Writing – original draft.

All authors read and approved the final manuscript.

## Acknowledgments

The authors thank the PSE-SANTE/SDOS/LMDN team of the IRSN for their excellent technical expertise on the MIRCOM facility and Valérie Buard (PSE-SANTE/SERAMED/LRMed) for her technical support on the IRSN confocal microscopy facility. The authors are grateful to P. Radicella for his critical reading of the manuscript, and to G. Gruel for his helpful suggestions.

## Funding

This work was supported by Institute National du Cancer, France (Grant number: PLBIO19-126 LS194750 INCa 2019).

## Data Availability Statement

The source data for the main and supplemental figures are available in the published article and its online supplemental files.

## Conflict of Interest Statement

The authors declare that they have no conflict of interest.

## Notes

### Competing Interest Statement

The authors have declared no competing interest.

### Summary of Updates

This version of the manuscript, has been revised for clarity of the text and additional results has been added as supporting data to enforce the discussion of the paper.

