## Supplemental Methods and Supplemental Figure Legends for "The DNA damage response in myogenic C2C7 cells depends on the characteristics of ionizing particles"

### Supplemental data

### Supplemental materials and methods

### Whole cell extracts and immunoblotting

We washed untransfected and stably GFP-tagged protein-expressing C2C7 myogenic cells, both under myoblast and myotube forms, twice in ice-cold PBS. Then, we immediately resuspended and lysed cells in Laemmli buffer containing 50 mM Tris pH 6.8, 2% SDS, 5% glycerol, 2 mM DTT, 2.5 mM EDTA, 2.5 mM EGTA, and 10 mM sodium fluoride (Sigma-Aldrich), supplemented with a cocktail of protease (Complete, EDTA-free tablets; Roche) and phosphatase (ThermoFisher Scientific) inhibitors, as described previously (Maire et al, 2013). Subsequently, we boiled cell lysates at 95 °C for 5 min and we then sheared them by syringing through a 26G needle. We quantified the protein concentration by using the BCA Protein Assay Kit-Reducing Agent Compatible (Pierce, ThermoFisher Scientific).

For each extract, we separated 15 µg of protein on a 4-12% TGX gels (BioRad) and transferred onto nitrocellulose membranes (BioRad). We saturated membranes with PBS-T containing 5% skimmed milk powder (Dutsher) and incubated them overnight at 4 °C with the appropriate primary antibodies diluted in TBS-T+ 5% milk. After washes, we incubated membranes with the appropriate HRP peroxidase secondary antibodies for one hour at RT. For visualization of proteins, we incubated membranes with Clarity ECL solution (BioRad) for chemiluminescence detection, imaged and analyzed membranes using the ChemiDoc MP Imaging System (BioRad) and ImageJ software (version 1.53e).

### Antibodies

Primary antibodies used during immunoblotting experiments are: rabbit anti-53BP1 (Bethyl, Euromedex, A300-272A; 1:1,000); mouse anti-β-actin (Sigma, A5441; 1:1,000) ), which was used as loading control; rabbit anti-H2AX (Cell Signaling Technologies, Ozyme, 2595; 1:1,000); mouse anti-γH2AX (Millipore, UpState, 05-636; 1:2,000); mouse anti-HP1α (Euromedex, 2HP-2G9-AS; 1:1,000); mouse anti- KU80 (Abcam, Ab119935; 1:200); mouse anti-myosine heavy chain (eBioSciences, ThermoFisher Scientific, MF-20, 14-6503-82; 1:500), which was used a marker of myotube state; rabbit anti-PARP1 (Enzo Life Sciences, ALX-210+302; 1:1,000); rabbit anti-RAD51 (Cell Signaling Technologies, Ozyme, 8875; 1:1,000) and β-tubulin (Sigma, TUB 2.1, T4026; 1:1,000), which was also used as loading control.

Secondary antibodies used are: horseradish peroxidase-conjugated affinity-purified donkey anti–mouse or anti–rabbit (1:5,000; Jackson ImmunoResearch Laboratories).

### Quantitative Image-Based Cell Cycle Analysis by Epifluorescence Microscopy

In order to determine how irradiated cells progress through the cell cycle, we reacquired the same samples (see “Immunostaining and image acquisition” paragraph of Materials and methods) on an inverted epifluorescence Olympus IX81 SCAN IM IX2 stage (Märzhäuser, Wetzlar, Germany), a MT20 fluorescence illumination microscope with a 10×/0.25 NA objective. The microscope is coupled with a motorized system with a fast filter wheel and an Orca R2 CCD camera (Hamamatsu, Massy, France). We adjusted the acquisition times for the different channels to obtain images under non-saturating conditions (i.e., in the 12-bit dynamic range) for all the experimental time-points analyzed. After acquisition, we performed image analysis with the ScanˆR analysis 4.2.1 software (Olympus, Rungis, France), as described previously (Toledo et al, 2013; Gruel et al, 2016). Briefly, we used an edge segmentation algorithm implemented in the software, which is based on Canny’s method to detect nuclei in the DAPI channel (main object) and γH2AX signal in the Texas Red channel (sub-object). We performed a first selection based on the area and circularity of the nuclei to consider only isolated nuclei and to remove from the analysis objects corresponding to clusters of nuclei and cellular debris. As in a flow-cytometry analysis, the cells were distributed in different phases of the cell cycle by assessing the integrated intensity of the DAPI signal (DNA content) within the entire nucleus. We visually determined the region containing the irradiated myoblasts through the γH2AX signal. Finally, we determined the cell cycle stage of all irradiated myoblasts together and non-irradiated myoblasts for each time-point upon irradiation to be able to analyze a minimum number of nuclei.

***Statistical analyses***

All data shown in this paper result from at least three independent experiments.

In order to investigate the impact of the experimental conditions on each monocellular fluorescence intensities, we performed a partial least squares discriminant analysis (PLS-DA) on the monocellular GFP-KU80 recruitment, RAD51, and γH2AX signals for the classification of groups defined by their type and number of particles as well as their time-points post-irradiation. The PLS-DA is a supervised dimension approach aiming to project a given dataset (here the fluorescence intensity matrices) on a subspace optimally separated according to a categorical variable (here the experimental conditions). Similarly, to its unsupervised dimension reduction counterpart, the well-known principal component analysis (PCA), the interpretation of PLS-DA consists of examining the contribution of the original covariates in the construction of its dimension components (called loadings).

### Supplemental figure legends

**Figure S1. Characterization of myogenic C2C7 cells expressing GFP-fused proteins.**

**A, B** Immunoblot analysis of GFP-tagged proteins produced by myogenic C2C7 cells stably transfected. Whole cell extracts of myogenic cells at the state of myoblast and myotube were probed with specific antibodies against 53BP1, KU80 and HP1α (upper panels) or with an anti-GFP antibody (bottom panels) (A) or against others DDR proteins, as PARP1, RAD51 and γH2AX (B). The positions of endogenous and GFP-tagged proteins are indicated in (A). * indicates in 53BP1-GFP-expressing myogenic cells the presence of truncated 53BP1-GFP proteins at a very low signal level. Actin and tubulin are used as loading control in (A, B) and myosin heavy chain as marker of myotube state in (B). The sizes of protein molecular weight markers (MW) are indicated in kDa. The immunoblots presented here are representative of 4 independent experiments.

**C** Quantification of protein expression levels from (A). We quantified and normalized the expression levels of endogenous and GFP-tagged proteins using ImageJ. Β-actin was used as loading control. Values represent the mean of at least 3 independent experiments and error-bars the standard deviation. No statistical significance was found by Student’s t test.

**Figure S2. Expression level of GFP-tagged proteins in myogenic C2C7 cells.**

**A-C** Representative images of myoblasts and myotubes stably expressing KU80-GFP (A), 53BP1-GFP (B) and HP1α-GFP (C). Borders of myotubes indicated by dashed white lines. Scale bar represents 10 µm.

**(D-F)** Quantification of GFP-tagged protein expression level by direct IF during myogenic differentiation, in C2C7 myoblasts (MB) and their committed progeny, myotubes (MT). The measured GFP fluorescence intensity of KU80-GFP (D), 53BP1-GFP (E) and HP1α-GFP (F) in myoblasts and myotubes are presented in scatter dot-plots, with bold bars representing the mean and error bars the standard error of the mean (SEM) from at least 120 nuclei/cell type for (D, E) and at least 59 nuclei/cell type for (F) and data are representative of 3independent experiments. Significance by unpaired t-Test, with ns for p> 0.05.

**Figure S3. Kinetic curves of KU80-GFP recruitment to the site of α-particle and proton micro-irradiation within myonuclei of differentiated myotubes.**

**A, B** Representative images of KU80-GFP recruitment in stably KU80-GFP-expressing C2C7 myotubes upon localized irradiation with reducing number of α-particles delivered (A) or protons (B). The localized irradiation site is marked with white arrowhead. Scale bar represents 10 µm.

**C, D** Kinetic curves of KU80-GFP recruitment obtained from A and B upon localized irradiation with a reducing number of α-particles (C) and protons (D). Error-bars represent the SEM obtained from N= 3 independent experiments with 46 - 85 nuclei / condition analyzed in (C) and 12 - 67 nuclei / condition analyzed in (D). The kinetic curves of KU80-GFP recruitments obtained in C2C7 myoblasts (Fig. 1) are also shown only for the irradiation with highest number of particles (i.e., 1,000 α-particles in (C) and 10,000 protons in (D)) by a very light blue curve. Statistical analysis performed by using generalized additive models (GAM) framework model and comparing each time point. Dashed boxes represent the region of slopes with significant difference between the kinetic curves. Color code of boxes represents the same color kinetic curve vs curve of highest number of particle irradiation. **** p< 0.00001.

**Figure S4. Accumulation of 53BP1 upon localized α-particle and proton irradiation in post-mitotic myotubes.**

**A** Recruitment kinetics of 53BP1-GFP in stably 53BP1-GFP-expressing C2C7 myotubes upon 1,000 and 50 α-particles or 10,000 protons irradiation. N= 3 independent experiments, mean ± SEM from 12 - 38 nuclei / condition analyzed. Statistical analysis performed by using GAM framework model and comparing each time point. Dashed boxes represent the region of slopes with significant difference between the kinetic curves. Color code of boxes represents the same color kinetic curve vs curve of highest number of particle irradiation. **** p< 0.00001.

**B** Representative images of stably KU80-GFP-expressing myoblasts (MB) and myotubes (MT) immunostained with antibodies against 53BP1 and γH2AX at the indicated time-points upon localized irradiation by 1,000 α-particles. DNA counterstained with DAPI. Scale bar, 10 µm.

**C** Relative fluorescence intensities at the site of localized irradiation of 53BP1 revealed by immunostaining at the indicated time upon targeted irradiation by 1,000 α-particles of stably KU80-GFP-expressing myoblasts and myotubes from (B). N= 3 independent experiments with 22 - 122 nuclei / condition analyzed. Data are presented in scatter dot-plots with bold bars representing the mean and error bars the SEM. Statistical significance is obtained by 2-way ANOVA with post-hoc Tukey’s multiple comparison test between myocytes and myotubes, with * p< 0.05, ** p< 0.01, *** p< 0.001.

**Figure S5. Balance between NHEJ and HR.**

**A, B** Mean nuclear fluorescence intensities of γH2AX staining (A) and, RAD51 staining and KU80-GFP signal (B) within nuclei of stably KU80-GFP-expressing C2C7 myoblasts at localized irradiation site 1 h after α-particles micro-irradiation with a reducing number of particles delivered. N= 3 independent experiments with 38 - 177 cells / condition analyzed. Data are presented in scatter dot-plots with bold bars representing the mean and error bars the SEM. Significance by 1-way (A) and 2-way (B) ANOVA with post-hoc Tukey’s multiple comparison test against the number of particles targeted. significant p-value are ns p> 0,05, * p< 0.05, ** p< 0.01, *** p< 0.001.

**C, D** Cell cycle distribution of non-irradiated neighboring myoblasts (C) and targeted myoblasts (D) upon irradiation with α-particles (Di) or protons (Dii), analyzed by immunofluorescence. Each value shown represents the mean of at least two independent irradiations. For each value, we analyzed in total between 80 and 800 nuclei. The error bars represent the standard error, and the p values are indicated (* p < 0.05).

**Figure S6. Diversification of C2C7 myoblasts depending on the fluorescence intensities of RAD51 and γH2AX staining and KU80-GFP signal upon micro-irradiation with α-particles targeted over time.**

**A, B** The partial least squares discriminant analysis (PLS-DA) scatterplots of data from Fig. S5 and Fig. 3, B-E according to the numbers of α-particles delivered (A), and according to the periods of time post-IR (B). The PLS-DA results in panels A-B were obtained from the α-particles dataset (986 cells). The significant contribution of RAD51, KU80 (KU80-GFP) and γH2AX intensities provided by the PLS-DA loading weight analysis are indicated with “+” symbol for each group.

**C, D** The position of each group of data from PLS-DA (A, B) are represented by density plots (in red) according to the numbers of α-particles delivered (C), and according to the periods of time post-IR (D).

**E** The performance measurement of the separation of each group of data to other conditions is estimated by the aera under the curves of the PLS-DA model. The AUC and the p value for each classification are shown in table.

**Figure S7. Diversification of C2C7 myoblasts depending on the fluorescence intensities of RAD51 and γH2AX staining and KU80-GFP signal upon micro-irradiation with protons targeted over time.**

**A, B** PLS-DA scatterplots of data upon irradiation with 10,000 protons, from Fig. 4, A-C, against α-particle irradiation with different number of particles from Fig. S6, A (A) and then over period of time post-IR from Fig. S6, B (B). The PLS-DA results in panels A-B were obtained from the α-particles and protons merged datasets (986 and 183 cells respectively). The significant contribution of RAD51, KU80 (KU80-GFP) and γH2AX intensities provided by the PLS-DA loading weight analysis are indicated with “+” symbol for each group.

**C, D** The position of each group of data from PLS-DA (A, B) are represented by density plots (in red) according to the numbers of α-particles or protons delivered (C), and according to the periods of time post-IR (D).

**E** The performance measurement of the separation of each group of data to other conditions is estimated by the aera under the curves of the PLS-DA model. The AUC and the p value for each classification are shown in table.
