## Supplemental Table 1 for "The DNA damage response in myogenic C2C7 cells depends on the characteristics of ionizing particles"

|  | **LET**  **(keV/µm)** | **Mean EDep (keV)** | **Nb particle** | **Mean Dose/dot (Gy)** | **Nb sDSB** | **Nb cDSB** |
| --- | --- | --- | --- | --- | --- | --- |
| **4 MeV proton** | 10 keV.µm^-1^ | 35,04 |  |  |  |  |
|  |  |  | **500** | 7.2 | **57.2** | **8.6** |
|  |  |  | **2,500** | 35.9 | **285.9** | **43.0** |
|  |  |  | **10,000** | 143.4 | **1143.8** | **172.1** |
| **6 MeV α-particle** | 84 keV.µm^-1^ | 346,2 |  |  |  |  |
|  |  |  | **50** | 7.1 | **109.7** | **71.9** |
|  |  |  | **200** | 28.3 | **438.9** | **287.6** |
|  |  |  | **1,000** | 141.7 | **2,194.3** | **1,438.2** |

**Table S1- Estimated number of simple DNA double strand breaks (sDSB) and complex DSB (cDSB, defined as 1 DSB with 1 or more strand breaks) for irradiations at a single dot within myoblast nuclei by defined number of 4 MeV proton or 6 MeV α-particle.**

These estimations were obtained from “dsbandrepair”, a Geant4-DNA simulation tool (Meylan et al, 2017; Tuan Anh et al, 2024) based on an extrapolation of experimental data from fibroblasts to those from myoblasts. This tool also estimates the mean energy deposit (Mean EDep) for each particle and the mean dose absorbed at the localized irradiation site.
