## Supplementary figures and images for "The DNA damage response in myogenic C2C7 cells depends on the characteristics of ionizing particles"

### Supplemental Figure 1

#### Sutcu et al., Figure S1

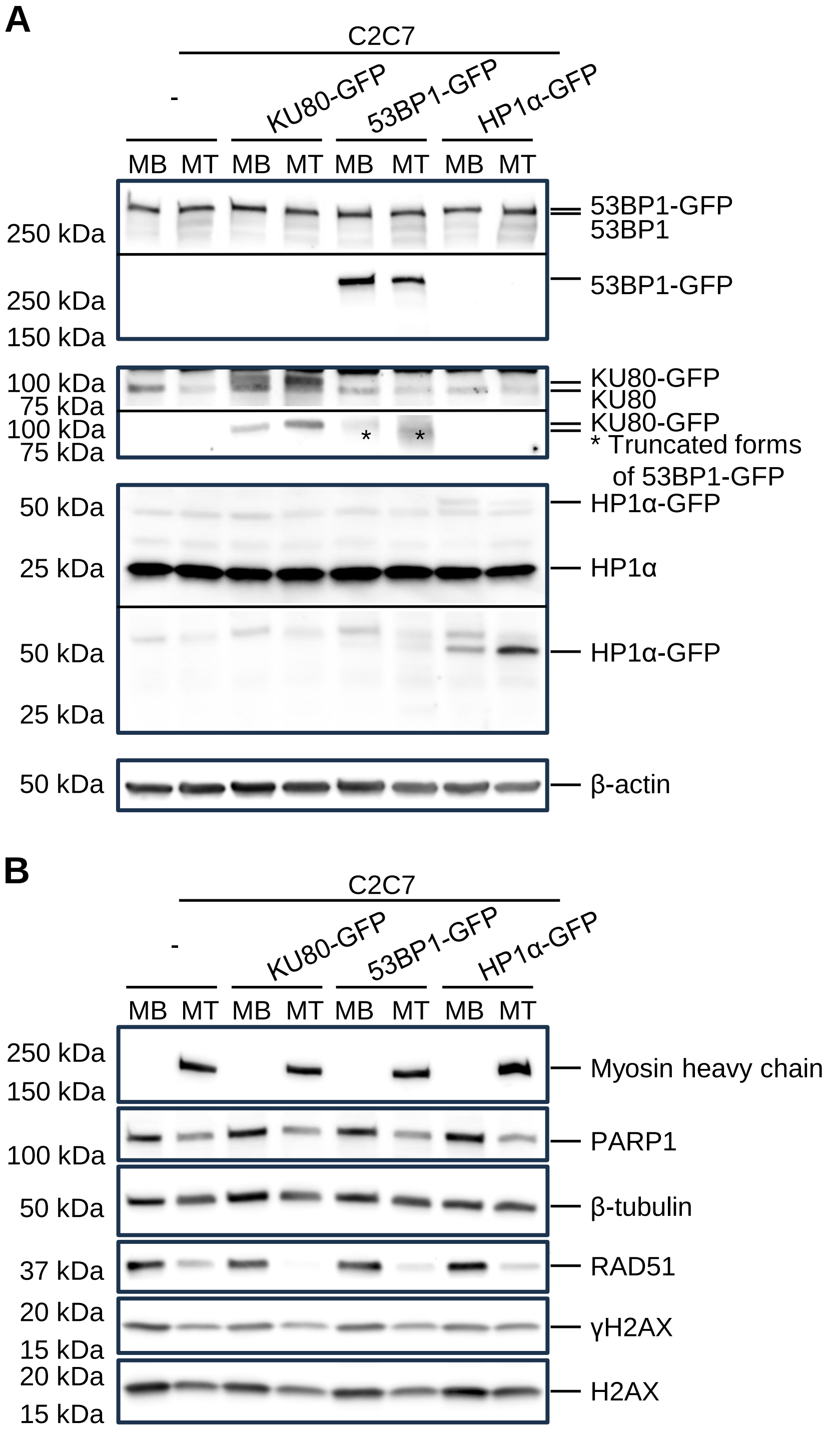


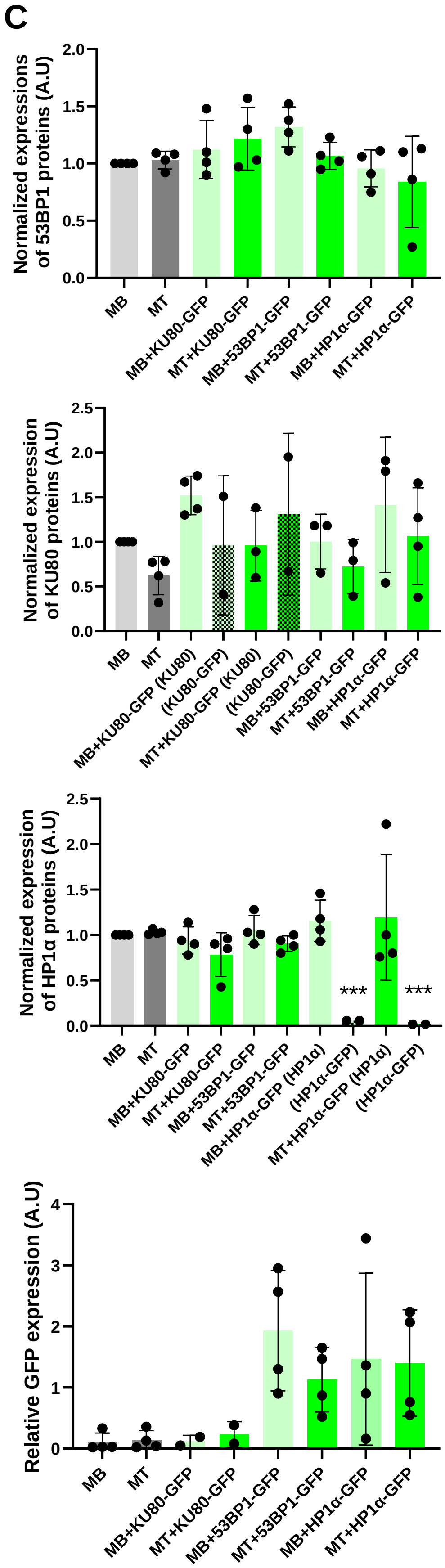

### Supplemental Figure 2

**Sutcu et al., Figure S2**

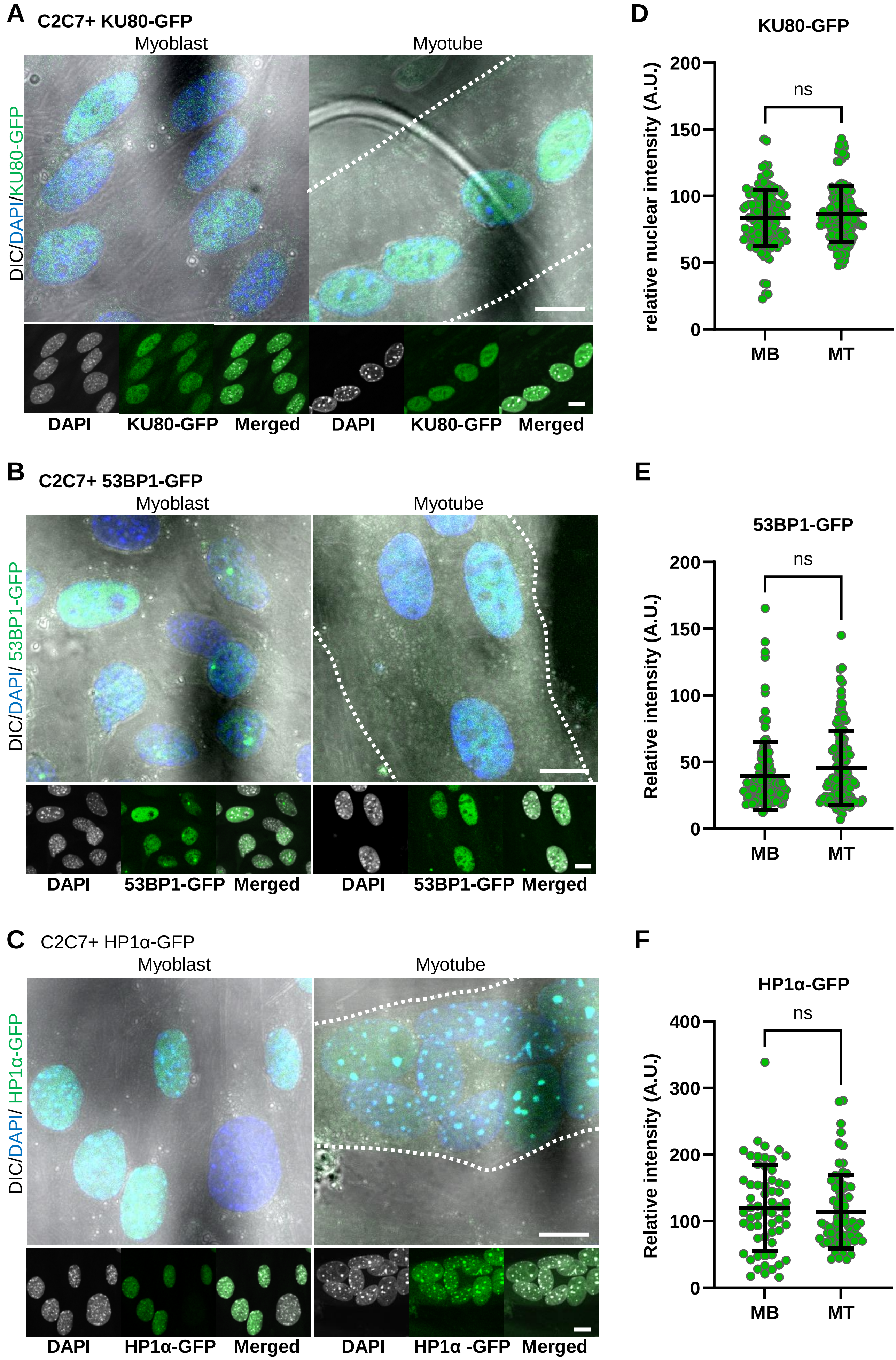

### Supplemental Figure 3

**Sutcu et al., Figure S3**

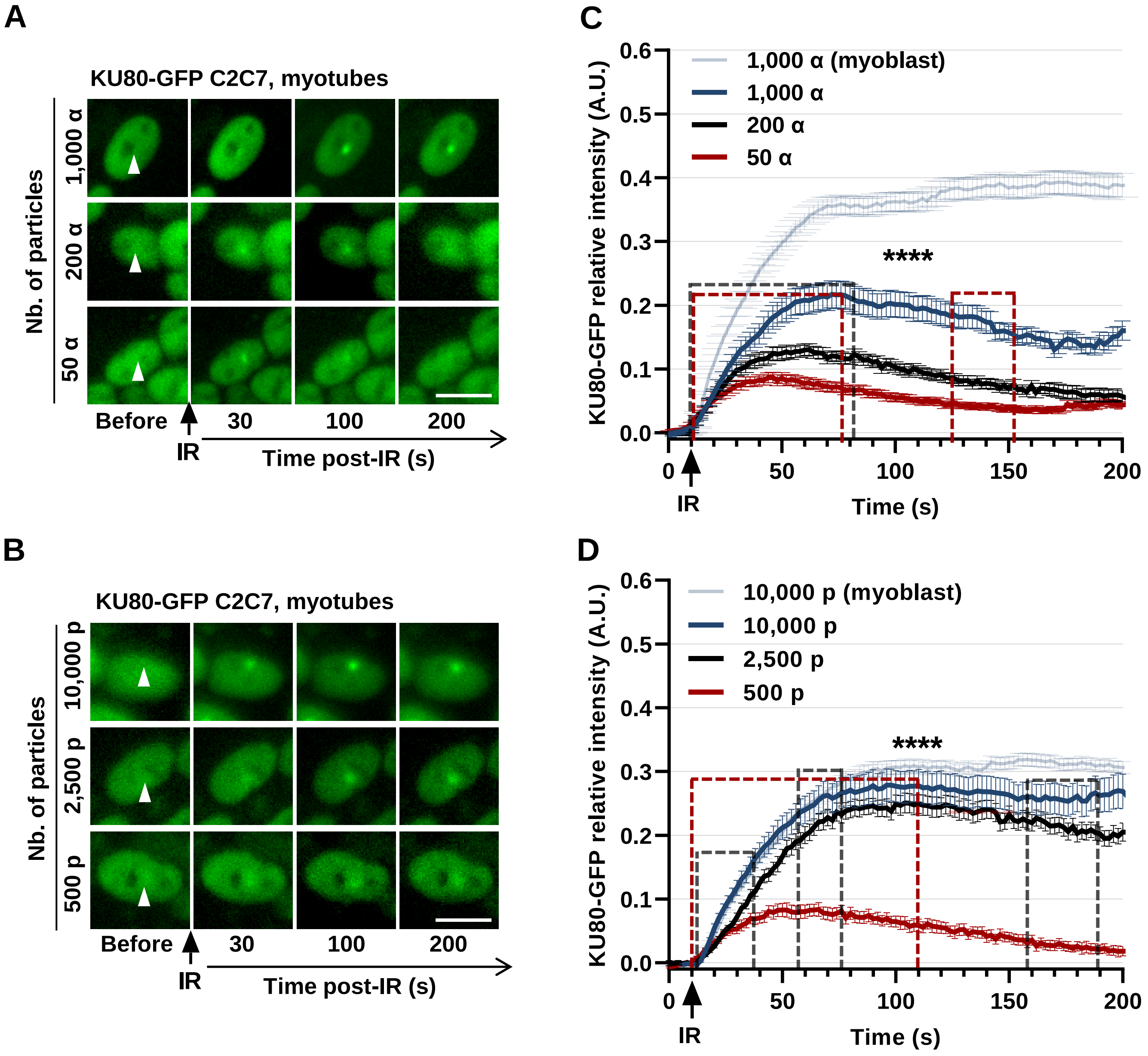

### Supplemental Figure 4

Sutcu et al., Figure S4

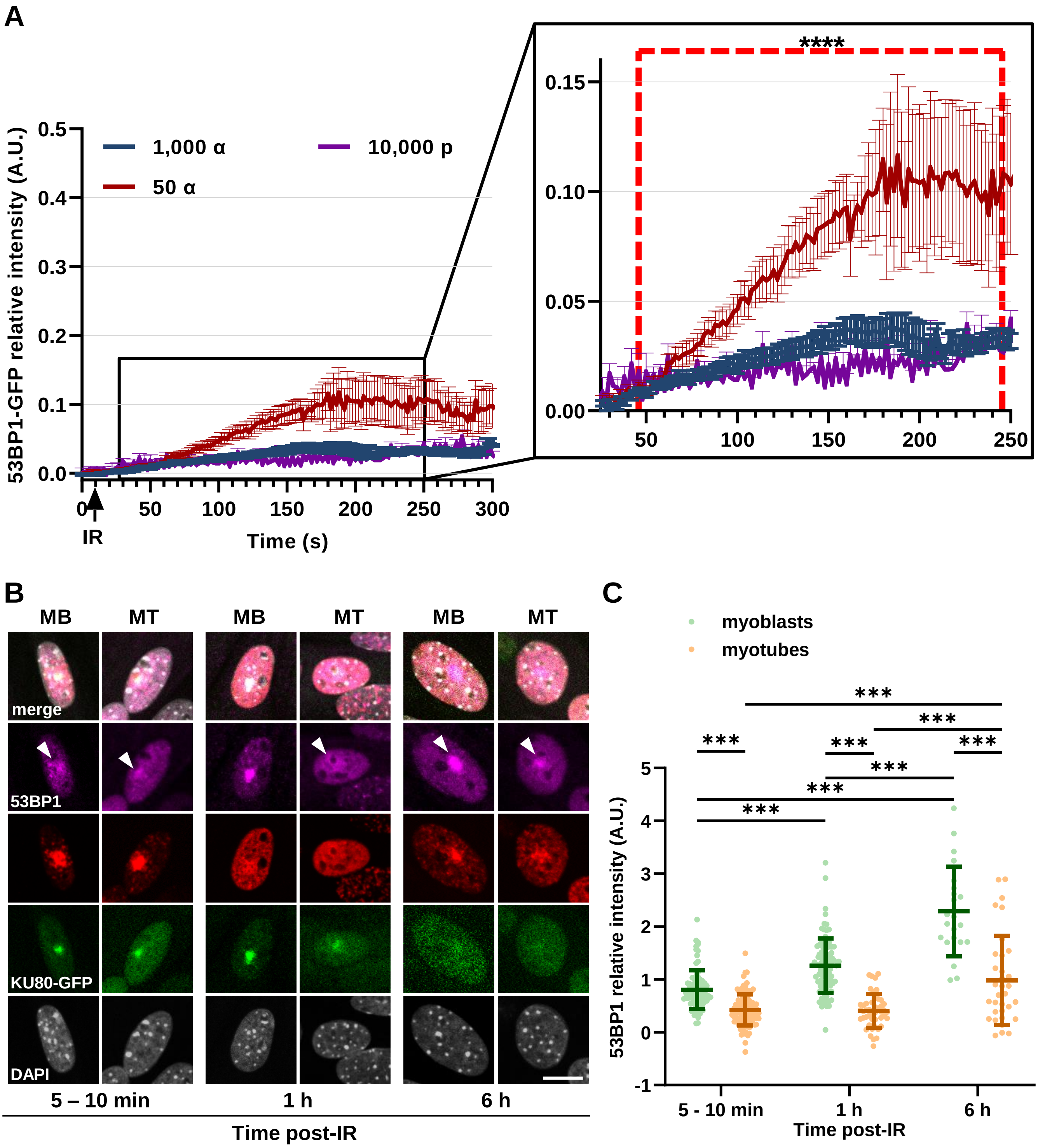

### Supplemental Figure 5

Sutcu et al., Figure S5

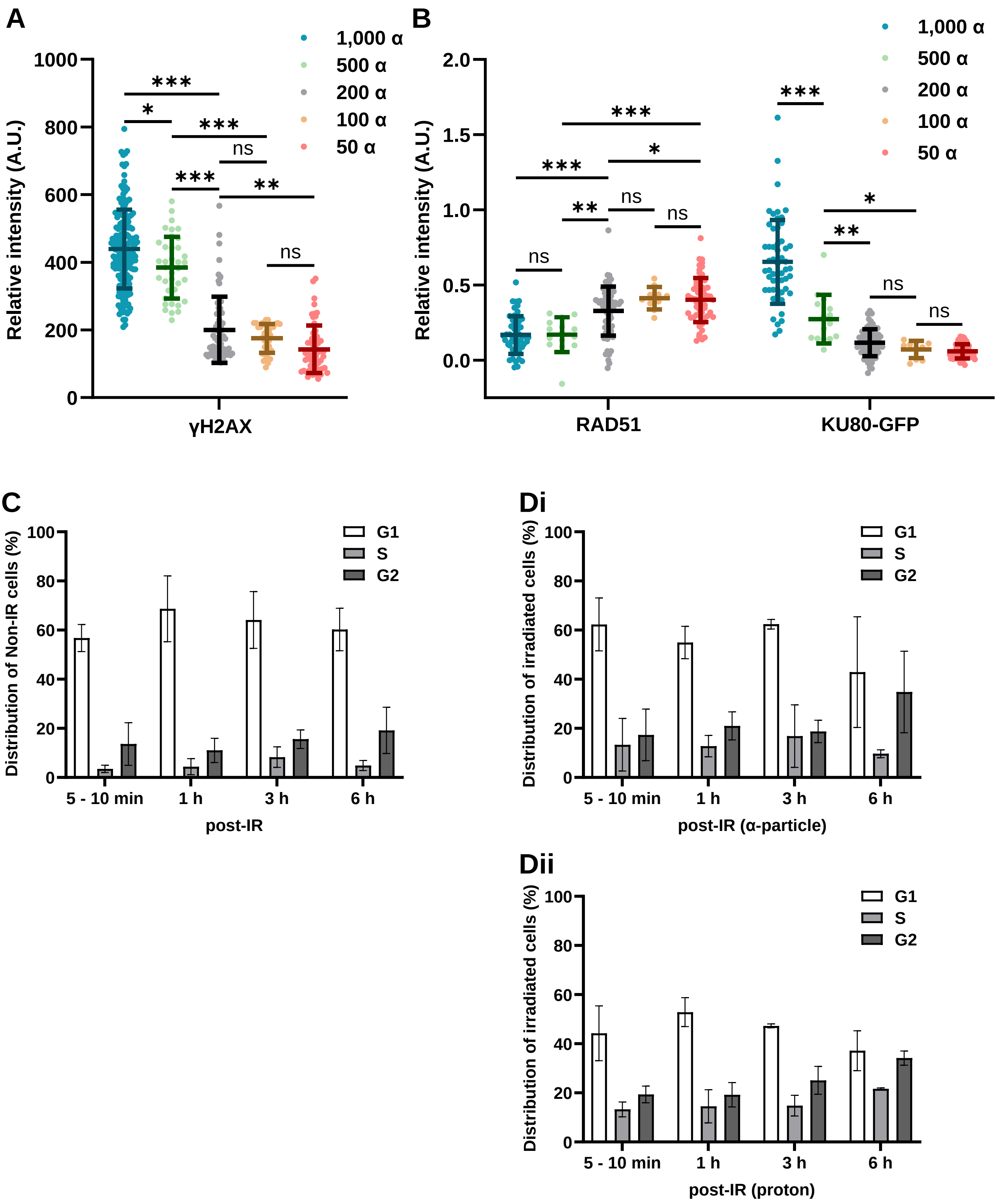
