## Supplemental Figure 6 for "The DNA damage response in myogenic C2C7 cells depends on the characteristics of ionizing particles"

### Sutcu et al., Figure S6

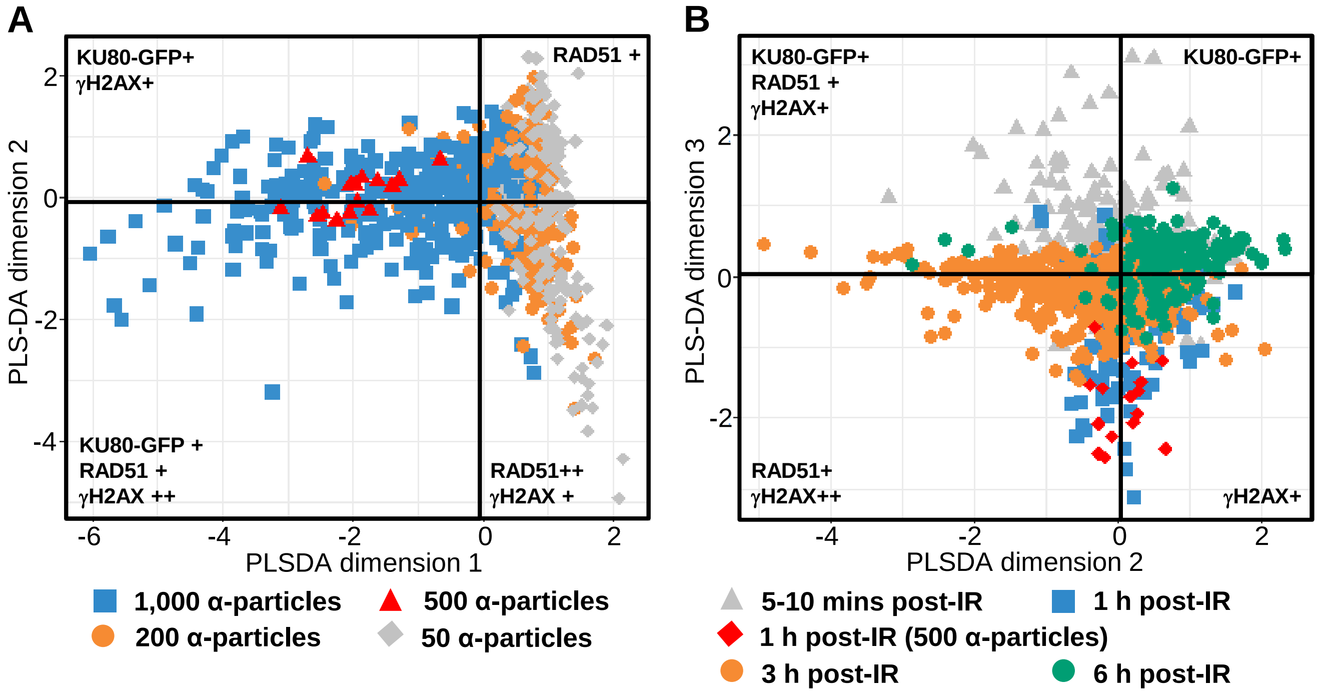


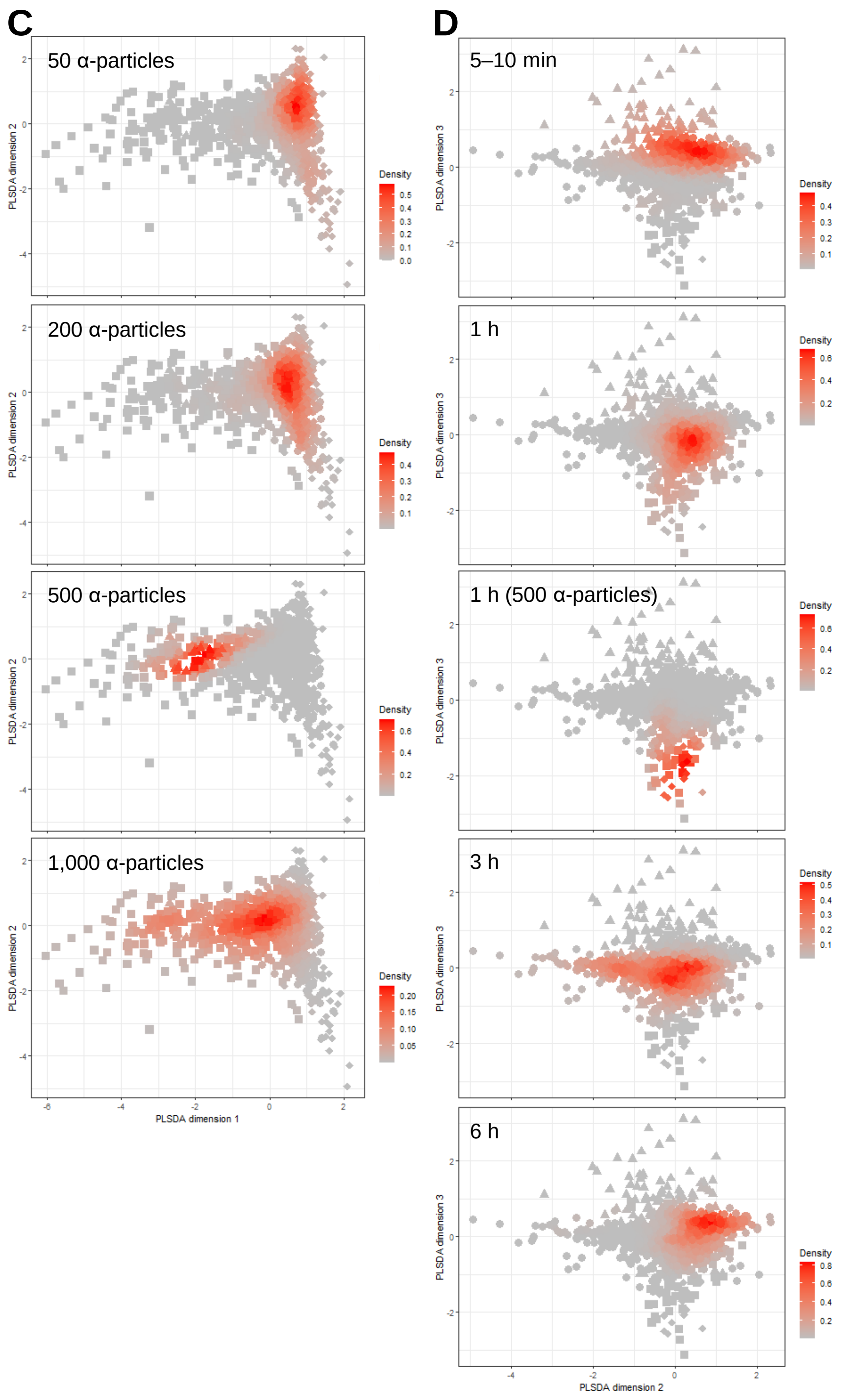


E

| | AUC | *p* value |
|---|---|---|
| 1,000 α_5-10 min vs others | 0.98 | 0 |
| 1,000 α_1h vs others | 0.95 | 0 |
| 1,000 α_3h vs others | 0.81 | 0 |
| 1,000 α_6h vs Others | 0.66 | $7.45^{e-06}$ |
| 200 α_5-10 min vs others | 0.76 | 2.2e-10 |
| 200 α_1h vs others | 0.77 | $1.11^{e-15}$ |
| 200 α_3h vs others | 0.77 | 0 |
| 200 α_6h vs others | 0.89 | 0 |
| 50 α_5-10 min vs others | 0.93 | 0 |
| 50 α_1h vs others | 0.76 | $1.01^{e-11}$ |
| 50 α_3h vs others | 0.79 | 0 |
| 50 α_6h vs others | 0.93 | 0 |
