## Supplemental Figure 7 for "The DNA damage response in myogenic C2C7 cells depends on the characteristics of ionizing particles"

### Sutcu et al., Figure S7

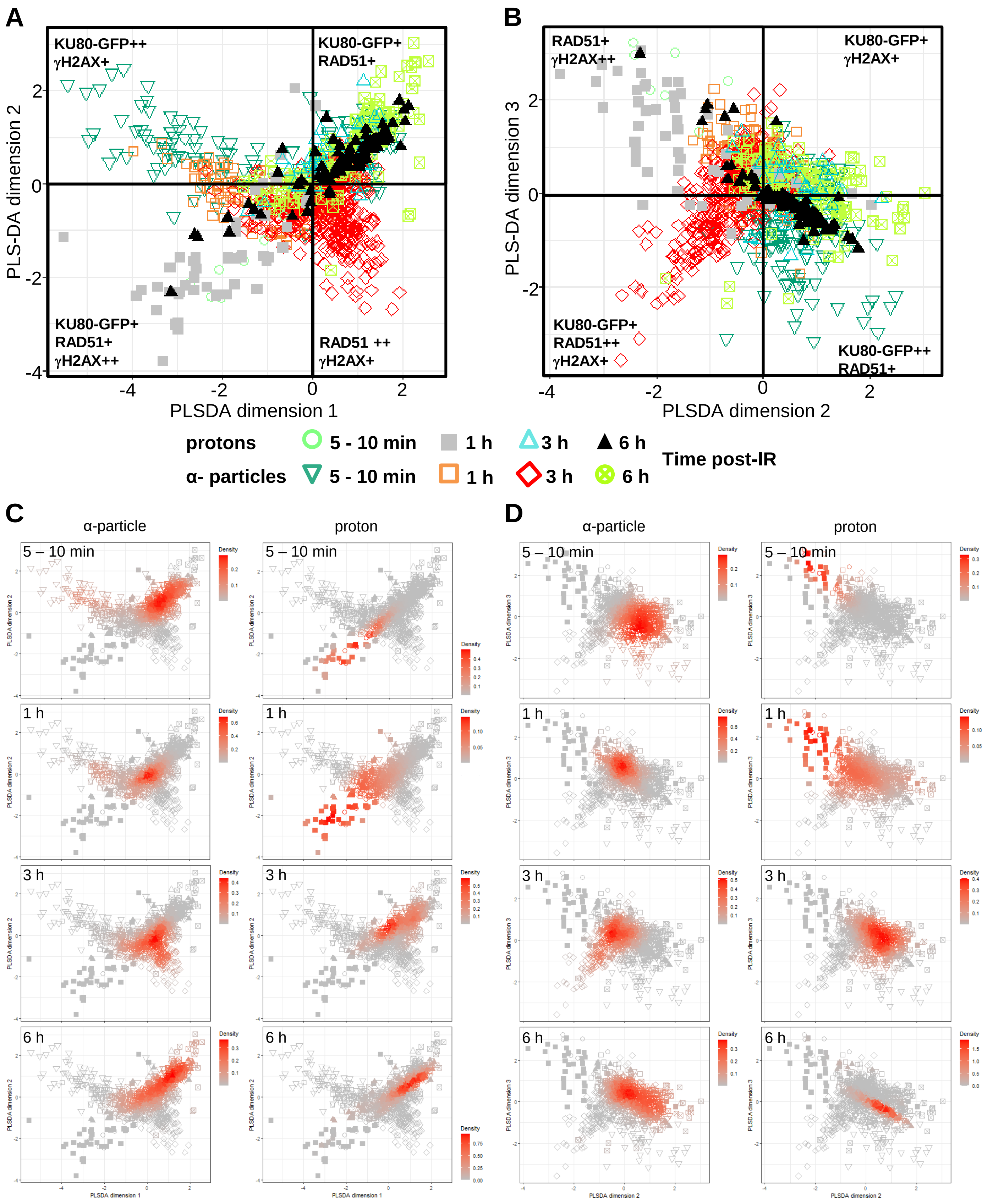


## E

| | AUC | *p* value |
|---|---|---|
| 50 α_5 min vs others | 0.93 | <0.001 |
| 50 α_1h vs others | 0.77 | <0.001 |
| 50 α_3h vs others | 0.79 | <0.001 |
| 50 α_6h vs others | 0.98 | <0.001 |

| | AUC | *p* value |
|---|---|---|
| 200 α_5 min vs others | 0.73 | <0.001 |
| 200 α_1h vs others | 0.81 | <0.001 |
| 200 α_3h vs others | 0.78 | <0.001 |
| 200 α_6h vs others | 0.89 | <0.001 |

| | AUC | *p* value |
|---|---|---|
| 1,000 α_5-10 min vs others | 0.97 | <0.001 |
| 1,000 α_1h vs others | 0.94 | <0.001 |
| 1,000 α_3h vs others | 0.84 | <0.001 |
| 1000 α_6h vs others | 0.71 | <0.001 |

| | AUC | *p* value |
|---|---|---|
| 10,000 Protons_10 min vs others | 0.9881 | <0.001 |
| 10,000 Protons_1h vs others | 0.871 | <0.001 |
| 10,000 Protons_3h vs others | 0.732 | <0.001 |
| 10000 Protons_6h vs others | 0.8074 | <0.001 |
| 3x 10,000 Protons_6h others | 0.8873 | <0.001 |
